# Ca^2+^-dependent vesicular and non-vesicular lipid transfer controls hypoosmotic plasma membrane expansion

**DOI:** 10.1101/2024.10.20.619261

**Authors:** Baicong Mu, David M. Rutkowski, Gianluca Grenci, Dimitrios Vavylonis, Dan Zhang

## Abstract

Robust coordination of surface and volume changes is critical for cell integrity. Few studies have elucidated the plasma membrane (PM) remodeling events during cell surface and volume alteration, especially regarding PM sensing and its subsequent rearrangements. Here, using fission yeast protoplasts, we reveal a Ca^2+^-dependent mechanism for membrane addition that ensures PM integrity and allows its expansion during acute hypoosmotic cell swelling. We show that MscS-like mechanosensitive channels activated by PM tension control extracellular Ca^2+^ influx, which triggers direct lipid transfer at endoplasmic reticulum (ER)-PM contact sites by conserved extended-synaptotagmins and accelerates exocytosis, enabling PM expansion necessary for osmotic equilibrium. Defects in any of these key events result in rapid protoplast rupture upon severe hypotonic shock. Our numerical simulations of hypoosmotic expansion further propose a cellular strategy that combines instantaneous non-vesicular lipid transfer with bulk exocytic membrane delivery to maintain PM integrity for dramatic cell surface/volume adaptation.

## Introduction

The plasma membrane (PM) forming the protective cell boundary, is a dynamic lipid bilayer interspersed with proteins, crucial for molecule exchange and stimulus response. Mechanically, lipid bilayers have a high elastic modulus, allowing only ∼ 3% pure elasticity-driven stretching under lytic tensions. ^1-3^ Therefore, to accommodate increasing cell volume and prevent cytolysis under conditions like hypoosmotic shock, excess membrane from surface and/or intracellular reservoirs must be incorporated to expand the PM surface area. Common membrane addition mechanisms, such as absorption of preexisting PM folds (e.g., eisosomes in yeast) and enhanced exocytosis seen in many cell types, ^4-6^ are thought to be regulated by membrane tension and calcium (Ca^2+^) signaling. ^6-10^ However, how mechanical sensing and subsequent membrane reorganization are coupled with the surface and volume adaptation process remains underexplored.

Hypoosmotically induced cell swelling is commonly used to study the fundamentals underlying adaptive surface and volume regulation. Mechanically gated channels are integral membrane proteins that undergo conformational changes induced by increased PM tension, such as that resulting from hypoosmotic shock. ^11^ Prokaryotic mechanosensitive channels of large and small conductance (MscL and MscS) are well-characterized stretch-activated osmotic release valves that protect bacteria from lysis during severe hypotonic shock. ^12^ MscS-like ion channels have been identified in fungi and plants but not in animals. Many of these eukaryotic counterparts are also functionally implicated in osmotic homeostasis of cells or organelles. ^13-15^ Here, we expand functions of MscS-like mechanosensitive channels to Ca^2+^ flux control that is essential for hypoosmotic swelling of the fission yeast *Schizosaccharomyces pombe* (*S. pombe*) protoplasts. We further revealed an unexpected role of Ca^2+^-dependent non-vesicular lipid transfer at endoplasmic reticulum (ER)-PM contact sites in sustaining PM integrity and allowing its massive expansion under acute hypoosmotic shock.

In *S. pombe*, ER-PM contacts are mainly established by ER-resident VAMP-associated proteins (VAPs) Scs2 and Scs22 via interactions with major anionic phospholipids (PLs) in the PM. ^16,17^ ER-anchored extended synaptotagmins (E-Syts), known as tricalbins in yeast, also assist ER-PM contact formation in both yeast and mammalian cells through Ca^2+^-dependent binding of PM acidic PLs via their tandem C2 domains. ^18-20^ Both yeast and mammalian E-Syts use their conserved synaptotagmin-like mitochondrial-lipid-binding protein (SMP) domains to non-selectively transfer glycerolipids at ER-PM contacts in a Ca^2+^-regulated manner. ^21-24^ However, the biological significance of E-Syt-mediated non-vesicular lipid transfer remains ambiguous, although recent studies have implicated them in maintaining PM integrity under heat stress. ^25,26^

In this study, we identified critical events underlying massive PM expansion upon acute hypoosmotic shock in *S. pombe* protoplasts. We also developed a mathematical model coupling major PM remodelling events with PM tension- and Ca^2+^-dependent regulatory mechanisms to depict cell expansion for osmotic equilibrium. Our study highlights E-Syts-mediated direct lipid transport as an emergency response to damage- or stress-induced Ca^2+^ influx to preserve PM integrity. Collectively, we propose that combining rapid non-vesicular lipid transfer with bulk exocytic membrane flux provides a robust mechanism for protecting PM integrity during drastic cell surface and volume adaptation.

## Results

### Extracellular Ca^2+^ influx controlled by MscS-like channels is essential for hypotonic PM expansion

To achieve osmotic equilibrium, cell swelling driven by hypoosmotic pressure requires coordinated expansion of the cell surface area. Inadequate PM addition increases its in-plane tension to lytic thresholds, leading to cell rupture. To investigate PM remodeling in response to severe osmolarity changes, we employed *S. pombe* protoplasts where cell walls were enzymatically removed (Figure 1A). Protoplasts were loaded into a 4.5 μm high microfluidic chamber perfused with steady-state isotonic buffer (see Materials and Methods) where they were slightly compressed and immobilized. An acute hypotonic buffer containing 0.2 M sorbitol was then introduced, creating an osmolarity change of ΔC = −1 M sorbitol (used for all experiments unless otherwise stated). Protoplasts were imaged for 10 minutes following the osmotic shift.

**Figure 1.**
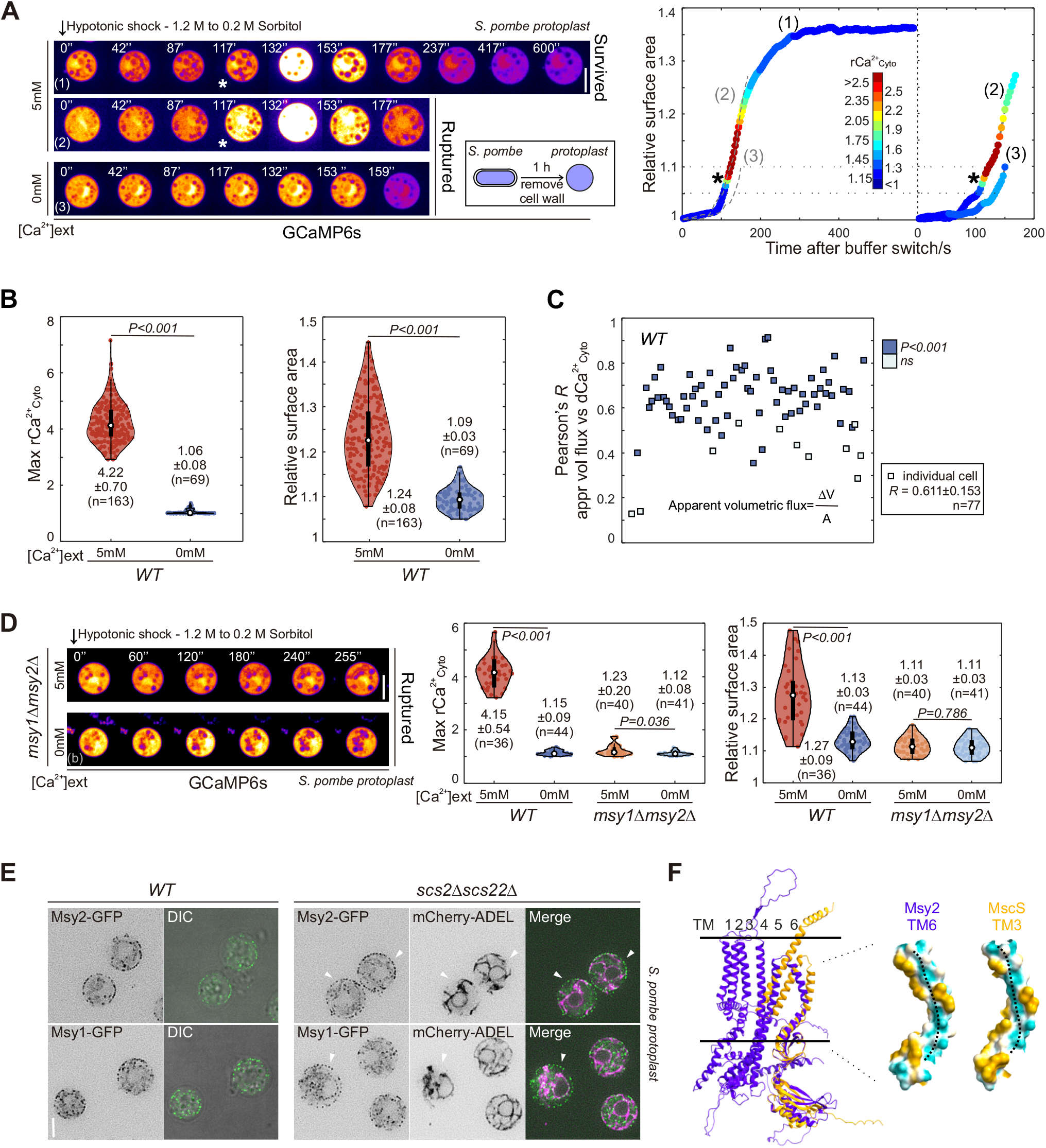
Extracellular Ca^2+^ influx controlled by MscS-like channels is essential for hypotonic PM expansion. (A and D) Time-lapse spinning disk confocal images of representative protoplasts expressing GCaMP6s with indicated hypotonic shocks. Shown are pseudo-colored images at central focal planes. Times, relative to buffer switch. Graph in (A) shows relative PM surface expansion curves of (1)-(3) protoplasts with colors denoting total relative cytosolic Ca^2+^ level (rCa^2+^_Cyto_). Dash lines mark expansion curves of indicated cells. Asterisks, the first time point of Ca^2+^ influx. (B and D) Quantifications of maximum rCa^2+^_Cyto_ and maximum relative surface area (mean ± standard deviation [SD]) of indicated protoplasts after 10-min acute hypotonic shock of indicated conditions. (C) Pearson correlation coefficients (R or Pearson’s R, mean ± SD) between apparent volumetric flux and Ca^2+^ influx rate (dCa^2+^). Each box represents an individual cell. n, cell number; *P*-values, two-tailed t-test. (E) Central focal plane spinning disk confocal images of indicated protoplasts. Arrows, GFP signals in ER-free PM regions. Scale bar, 5 μm. (F) Structural alignment of AlphaFold-predicted monomeric Msy2 with MscS. Lining hydrophobicity map of TM6 (Msy2) and TM3 (MscS) is indicated with dotted lines showing putative water paths. Dark goldenrod and dark cyan denote the most hydrophobic and hydrophilic areas, respectively.

Interestingly, under acute hypoosmotic shock, the cytosolic Ca^2+^_Cyto_ level indicated by the GCaMP6s reporter^27^ (the relative total intensity is quantified as rCa^2+^_Cyto_, see Methods) dramatically increased during the rapid and massive expansion of wild-type (*WT*) protoplasts (Figure 1A). For surviving cells, the cytosolic Ca^2+^ level eventually decreased (Figure 1A). On average, the PM surface area increased 24% in *WT* protoplasts after acute hypotonic shock (Figure 1B). Notably, protoplasts exposed to the same osmolarity change in Ca^2+^-free buffer showed no significant increase in rCa^2+^_Cyto_ or PM expansion and were rapidly lysed after expanding around 10% of their initial surface area (Figures 1A and 1B). These data suggest that external Ca^2+^ is critical for substantial PM expansion following acute hypoosmotic shock, implicating Ca^2+^-dependent mechanisms in promoting PM integrity and/or PM addition.

We noticed that Ca^2+^ influx rate (dCa^2+^_Cyto_) was highly correlated with the apparent volumetric flux (defined as dV/A) during Ca^2+^ elevation (Figure 1C), indicative of coupled Ca^2+^-water inflow probably via channels. It is known that the PM tension surges during osmotic swelling, likely implying the involvement of mechanosensitive/stretch-activated ion channels in this process. Remarkably, protoplasts lacking both MscS-like mechanosensitive channels (Msy1 and Msy2) showed no apparent Ca^2+^ influx after hypoosmotic shock regardless of Ca^2+^ in the medium (Figure 1D). Among the two, Msy2 played a more dominant role in controlling such Ca^2+^ influx (Figure S1A). Like *WT* protoplasts treated with Ca^2+^-free hypotonic buffer, both *msy1*Δ*msy2*Δ and *msy2*Δ cells exhibited limited PM expansion (∼10%) before quick rupturing, independently of external Ca^2+^ (Figures 1D and S1A).

In fact, unlike a previous study that showed their ER localization via immunostaining, ^13^ C-terminally GFP-tagged Msy1 and Msy2 expressed from own native loci, mainly localized to the cell cortex in both walled cells and protoplasts (Figures S1B and 1E). Using *scs2*Δ*scs22Δ* background where the cortical ER (cER) indicated by the luminal ER marker mCherry-ADEL^28^ is largely detached from the PM, ^16^ we confirmed that a considerable portion of Msy2 and Msy1 localized to the PM of the protoplasts (Figure 1E), supporting their possible functions in gating Ca^2+^ influx in the PM during hypotonic shock. Intriguingly, unlike Msy2, Msy1 localized more to intracellular compartments in walled cells, including the ER, (Figure S1B), suggesting its gradual relocation to the PM during protoplast formation.

MscS-like channels facilitate ion flows along concentration gradients. Of note, rCa^2+^_Cyto_ always clearly dropped right before the cell rupture under Ca^2+^-free hypoosmotic shock in *WT* and *msy1Δ* protoplasts (Figures 1A and S1A). Such Ca^2+^ efflux might result from Msy2-mediated channel opening before the stretched PM reaches lytic tension. In line with this idea, such Ca^2+^ efflux was never seen in protoplasts lacking Msy2 (Figures 1D and S1A). Moreover, none of other putative stretch-activated ion channels^29^ or organellar Ca^2+^-ATPase^30^ appeared to be crucial for controlling Ca^2+^ flux or for hypotonic protoplast expansion (Figure S1C). Curiously, protoplasts lacking the only putative water channel Aqp1 exhibited normal Ca^2+^ influx and *WT-*like hypoosmotic PM expansion (Figure S1C). This seems to align with the previous notion that *S. pombe* lacks a bona fide water channel. ^31^ Given that the lining of Msy2 channels contains polar residues arranged similarly to those of the bacterial MscS heptamers (Figure 1F), ^32^ it may also mediate water inflow like MscS during hypotonic cell swelling.

Taken together, our results suggest that extracellular Ca^2+^ influx controlled by MscS-like mechanosensitive channels is essential for hypoosmotic PM expansion.

### Exocytosis and E-Syts are required for massive hypotonic PM expansion

We next investigated membrane sources for such massive hypotonic PM expansion. Exocytosis is known to increase the PM surface area via exocytic vesicle fusion. To confirm the role of exocytosis in massive hypotonic PM expansion, we pretreated protoplasts with Brefeldin A (BFA) which blocks ER-to-Golgi trafficking and thus ultimately prohibits exocytosis. ^33^ As expected, the PM expansion capacity of BFA-treated protoplasts was clearly decreased during osmolarity equilibrium after acute hypotonic shock, with the surface area increment before cell rupture dropping from 25% to 9% as pretreatment duration increased from 0 to 90 minutes (Figures 2A and 2B). Notably, 90-min BFA treatment that presumably exhausted existing exocytic vesicle pool before hypotonic shock, resulted in limited PM expansion (∼10%) before quick rupturing, similar to cells shocked with Ca^2+^-free buffer or *msy2*Δ mutants lacking Ca^2+^ influx after the shock (Figures 2B, 1B, 1D and S1A). These data suggest that exocytosis is essential for extensive hypotonic PM expansion and also imply a possible stimulatory role of elevated Ca^2+^ levels in exocytosis during hypotonic shock.

**Figure 2.**
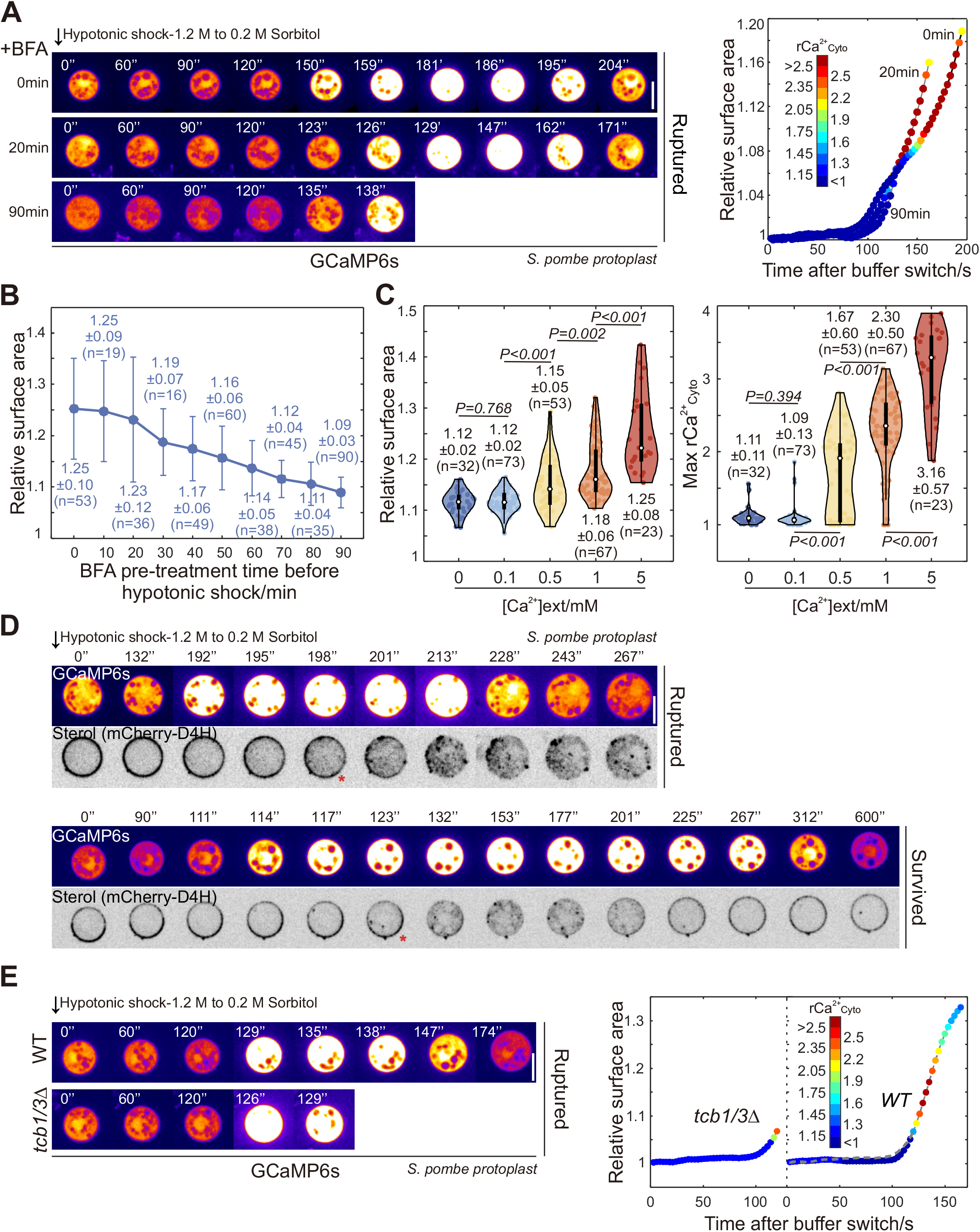
Exocytosis and E-Syts are required for massive hypotonic PM expansion. (A and E) Time-lapse spinning disk confocal images of representative protoplasts expressing GCaMP6s with indicated hypotonic shocks. Shown are pseudo-colored images at central focal planes. In (A), Duration of BFA pretreatment is indicated on the left. Graphs on the right show relative PM surface expansion curves of corresponding protoplasts with colors denoting rCa^2+^_Cyto_. The grey dash line in (E) mark expansion curves of *tcb1*Δ*tcb3*Δ. (B) Quantifications of maximum relative surface area (mean ± SD) of BFA*-*pretreated *WT* protoplasts after 10-min acute hypotonic shock. Error bars, 2×SD. (C) Quantifications of maximum rCa^2+^_Cyto_ and maximum relative surface area (mean ± SD) of *WT* protoplasts after 10-min hypotonic shock of indicated conditions. n, cell number; *P*-values, two-tailed t-test. (D) Time-lapse spinning disk confocal images of representative protoplasts expressing both GCaMP6s and mCherry-D4H. Shown are pseudo-colored GCaMP6s and inverted mCherry-D4H images at central focal planes. Times, relative to buffer switch. Scale bar, 5 μm.

The PM invaginations, known as eisosomes, have been proposed to be responsive to PM tension variation and act as membrane reservoirs. ^5,34^ Indeed, we also observed eisosome disassembly after acute hypotonic shock, primarily during massive PM expansion (Figure S2A). However, unlike the previous study showing limited expansion of eisosome-deficient *pil1*Δ protoplasts after hypoosmotic shocks, ^5^ we found *pil1Δ* protoplasts exhibited *WT*-like hypotonic PM expansion, with only a minor reduction in cell surface area increment (Figure S2B). Similarly, when exocytosis was pre-blocked, *pil1Δ* protoplasts showed slightly less PM expansion than *WT* after the shock (Figure S2C). These results suggest that eisosomal furrows play a minor role in hypotonic PM expansion.

One the other hand, endocytosis reduces the PM surface area. A brief pretreatment (∼10 mins) with Latrunculin A (LatA) that inhibits actin polymerization and thus endocytosis, ^35,36^ led to an apparent delay in Ca^2+^ influx after the shock (Figure S2D, see Materials and Methods). Conceivably, the lack of endocytosis may attenuate PM tension increase which is typically associated with a reduction in PM surface area, therefore likely delaying the activation of MscS-like channels responsible for Ca^2+^ influx. However, the overall hypotonic PM expansion of LatA-treated protoplasts remained *WT*-like (Figure S2B).

Exocytosis was shown to be accelerated in *S. pombe* protoplasts upon hypoosmotic shock. ^37^ However, unlike in neurons, the synaptotagmin-based mechanism of Ca^2+^-stimulated exocytosis^38^ is absent in fission yeast which lacks synaptotagmins. Moreover, both Ync13 and Git1, the *S. pombe* homologs of Munc13 family which is one core Ca^2+^ effector for synaptic vesicle priming, ^39^ were dispensable for massive hypotonic PM expansion (Figure S2E). Our results somewhat align with previous studies showing that none of them are involved in exocytosis. ^40,41^ Other potential Ca^2+^-effectors, including putative Rab GTPases Gyp2 and Gyp4 (encoded by SPBC215.01), which are implicated in vesicle trafficking, were also not required for hypotonic PM expansion (Figure S2E). Therefore, *S. pombe* appears to lack Ca^2+^-sensors that can immediately boost exocytosis. Curiously, Ca^2+^ influx-triggered massive PM expansion occurred only when Ca^2+^ concentration in the hypoosmotic buffer exceeded 100 μM (Figure 2C). However, constitutive exocytosis for fission yeast growth takes place at much lower Ca^2+^ levels of 100-200 nM. ^29^ We thus hypothesized that other mechanisms dependent on higher Ca^2+^ levels are involved in supporting hypotonic PM expansion.

We noticed that most BFA-treated protoplasts ruptured immediately after Ca^2+^ influx under acute hypotonic shock (Figure 2A), implying that the PM becomes vulnerable due to presumably further tension increase following activation of MscS-like channels. Indeed, PM integrity appeared compromised over this critical period, as indicated by a quick drop in the PM sterol levels shortly after Ca^2+^ influx (Figure 2D). We did not observe similar sudden level changes with other major signaling lipids in the PM at this injury juncture (Figure S2F). For protoplasts that survived the severe shock, PM sterol levels eventually restored (Figure 2D), implying that active repair mechanisms may help heal small PM wounds. Such hypothetical PM repair does not require Ca^2+^-dependent ESCRT-III/Vps4 machinery^42^, as no apparent defects were seen in hypotonic PM expansion in protoplasts lacking related key players (Figure S2E). Instead, our screening for Ca^2+^-effectors involved in hypotonic PM expansion revealed that *tcb1Δtcb3Δ* protoplasts lacking two fission yeast E-Syts barely survived such PM injury and ruptured quickly after Ca^2+^ influx with limited PM expansion (Figure 2E). Budding yeast E-Syts have been proposed to function in maintaining PM integrity. ^25,26^ Remarkably, efficient E-Syt-mediated non-vesicular lipid transfer is only activated by higher Ca^2+^ levels above 100 μM, ^22,24^ implicating E-Syts in this hidden mechanism proposed above.

In summary, we conclude that both exocytosis and E-Syts are essential for extensive hypoosmotic PM expansion, possibly with exocytosis mainly providing membrane reservoirs and E-Syts keeping PM integrity.

### E-Syts-mediated lipid transfer at ER-PM contacts is critical for hypotonic PM expansion

There are three E-Syt homologs in *S. pombe* (Tcb1-3), with each possessing the transmembrane (TM) segment anchoring it to the ER, one SMP domain with lipid transfer function, and multiple C2 domains that bind Ca^2+^ (Figure 3A). Among them, Tcb2 is functionally unrelated, as *tcb2Δ* protoplasts exhibited normal Ca^2+^-dependent hypotonic PM expansion (Figure 3B). In contrast, both *tcb1Δ* and *tcb3Δ* single mutants showed compromised PM expansion similar to *tcb1Δtcb3Δ* double mutant under acute hypoosmotic shock (Figure 3B), implying that they may function together. Unlike Tcb2 which was localized throughout ER compartments, Tcb1 and Tcb3 were mainly enriched at ER-PM contacts in both protoplasts and walled cells. These localization patterns were more evident in *scs2Δscs22Δ* background (Figures 3C and S3A). Interestingly, the accumulation of Tcb1 and Tcb3 at ER-PM contacts is mutually dependent: Tcb1 delocalized to the perinuclear ER in *tcb1Δ*, while Tcb1 levels in the cortical ER decreased in *tcb3Δ*. Their localization remained unchanged in *tcb2Δ* (Figures 3C and S3A). E-Syts were proposed to function as dimers. ^21,43,44^ Indeed, we confirmed Tcb1-Tcb3 interaction using co-immunoprecipitation and split-YFP-based bimolecular fluorescence complementation (Figures 3D and S3B). Collectively, our data suggest that Tcb1 and Tcb3 likely act as complexes at ER-PM contacts to facilitate hypotonic PM expansion.

**Figure 3.**
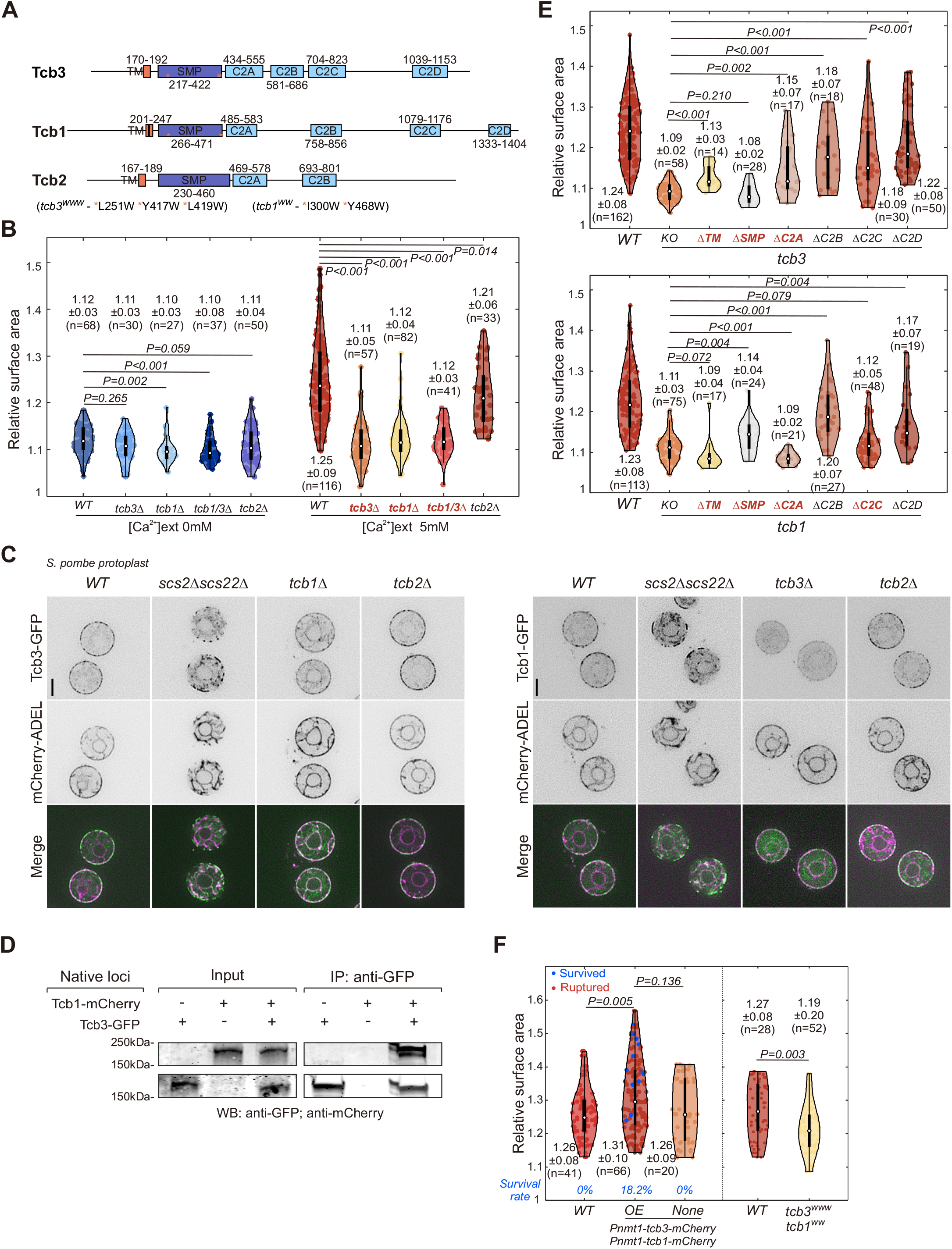
E-Syts-mediated lipid transfer at ER-PM contacts is critical for hypotonic PM expansion. (A) Schematic diagram of functional domains of three tricalbins. (B, E and F) Quantifications of maximum relative surface area (mean ± SD) of indicated protoplasts after 10-min hypotonic shock of indicated conditions. n, cell number; *P*-values, two-tailed t-test. In (F), survival rates are included; OE, overexpression. (C) Central focal plane spinning disk confocal images of indicated protoplasts. Scale bar, 5 μm. (D) Co-IP of indicated proteins expressed from native loci. Samples were probed with indicated antibodies.

Furthermore, our domain analysis using truncation mutants manifest that the TM, SMP and C2A domains of either E-Syts, as well as the C2C domain of Tcb1, are crucial for hypotonic PM expansion (Figure 3E and S3C). It has been shown that Ca^2+^-C2A interaction activates the adjacent SMP domain for lipid transfer, while Ca^2+^-C2C association facilitates membrane tethering. ^45^ Therefore, our results suggest that the ER-PM contact localization, lipid transfer activity and membrane tethering capacity of both E-Syts are essential for hypotonic PM expansion. In particular, truncation of the SMP domain in Tcb3 or the C2A domain in Tcb1 led to a complete loss of function (Figure 3E), implying that they may have prioritized roles - Tcb1 primarily in tethering and Tcb3 mainly in lipid transfer at ER-PM contacts.

To further substantiate that roles of these two E-Syts in PM expansion are pertinent to their lipid transfer function, we mutated several residues in the lipid-binding groove of their respective SMP domain to bulky tryptophan (W) to hamper lipid access through steric effects (Figure 3A). These mutations did not affect protein localization but indeed led to compromised hypotonic PM expansion (Figures 3F and S3C). Of note, the resultant mutant *tcb1*^*ww*^*tcb3*^*www*^ was less sensitive than mutants with complete loss of Tcb1 or Tcb3 function, likely due to their residual lipid transfer activities, as the similarly mutated mammalian E-Syt1 was also not fully inactive. ^22^ Conceivably, small PM wound healing *per se* may not require much lipid addition which *tcb1*^*ww*^*tcb3*^*www*^ could still provide. The reduced PM surface increase in *tcb1*^*ww*^*tcb3*^*www*^ background implies that Tcb1/3-mediated lipid transfer may also contribute to expanding the PM. In line with this, raising the amount of Tcb1/3 resulted in a ∼5% more PM surface area expansion and a ∼18.2% higher survival rate during the shock (Figures 3F and S3D).

Surprisingly, despite Tcb1/3-mediated lipid transfer occurring at ER-PM contact sites, hypotonic PM expansion of *scs2Δscs22Δ* protoplasts which lack these contacts, remained *WT-* like (Figures S3E and 3C). It has been shown that exocytic rate is higher in *scs2Δscs22Δ* cells due to increased PM availability for vesicle fusion. ^46^ We thus reasoned that any deficit in hypotonic PM expansion of *scs2Δscs22Δ* due to decreased Tcb1/3 levels at ER-PM contacts might be offset by enhanced exocytosis. Indeed, when exocytosis was pre-inhibited by BFA, PM surface increment was reduced more in *scs2Δscs22Δ* than *WT* after the shock (Figure S3E). When exocytic vesicle pool was exhausted by 90-min BFA pretreatment before the shock, both protoplasts exhibited similarly limited PM expansion (Figure S3E), again suggesting that exocytosis is the major membrane source for PM surface increase.

Collectively, we propose that Tcb1-Tcb3 complexes maintain PM integrity and supply lipids required for PM surface expansion via Ca^2+^-activated lipid transfer at ER-PM contacts during hypoosmotic shock.

### Mechanistic depiction of hypotonic PM expansion by a numerical model

Combining the key cellular events hitherto identified with principles of membrane mechanics, we developed a numerical model to describe the hypotonic PM expansion process for osmotic equilibrium. This model primarily focused on the coordinated remodeling events of various membrane sources (i.e., endo- and exocytosis, eisosome turnover, and non-vesicular lipid transfer), regulated by PM tension and/or Ca^2+^, to allow cell surface expansion to accommodate water flux driven by osmotic and hydrostatic pressure differences (Figure 4A, see Materials and Methods). For simplicity, we treated the PM as a single mechanical structure and neglected the contributions of cytoskeletal mechanics to PM tension variation.

**Figure 4.**
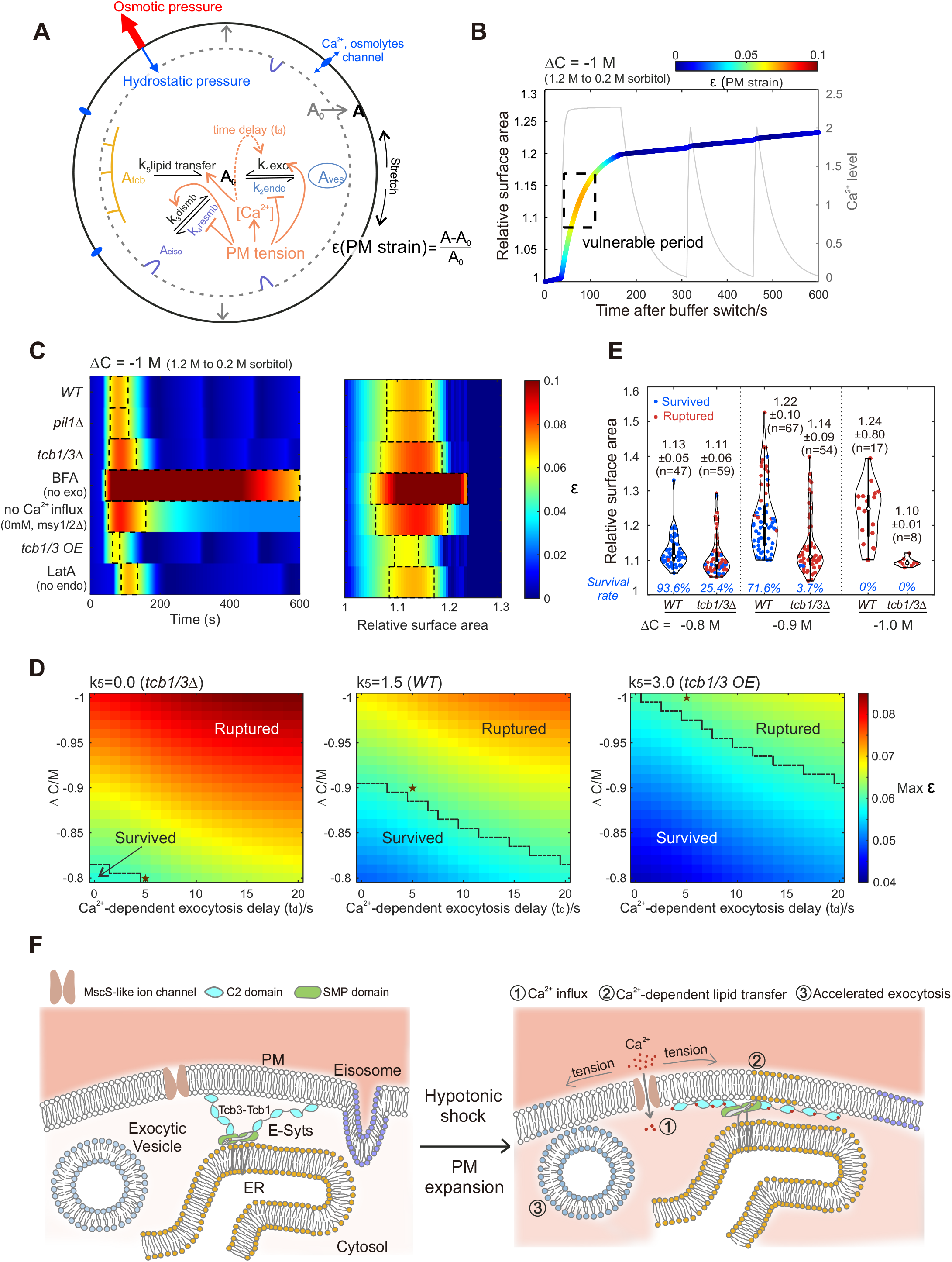
Mechanistic depiction of hypotonic PM expansion by a numerical model. (A) Schematic diagram of the mechanistic model. A_0_, unstretched PM area; A_ves_, area of exocytic vesicle pool; A_tcb_, membrane area from E-Syts-mediated lipid delivery; A_eiso_, eisosomal membrane area; k_1_ to k_5_, rates of exocytosis, endocytosis, eisosome disassembly, eisosome reformation and lipid transfer. [Ca^2+^], cytosolic Ca^2+^ elevation. (B) Simulation of relative PM surface area expansion with colors indicating PM strains (ε) during 10-min hypotonic expansion. The framed vulnerable period is defined as ε above 0.06. The corresponding Ca^2+^ variation (grey) curve is included. (C) Plots of simulated ε variation in indicated model conditions along hypoosmotic expansion time (left) and relative surface area (right). Vulnerable periods are framed. (D) Maximum ε in simulations of different Ca^2+^-dependent exocytosis delays versus varying osmolarity differences. Ruptured or survived zones are defined as the maximum ε above or below 0.06. Stars, predicted scenarios at t_d_=5 s that match experimental results. (E) Quantifications of maximum relative surface area (mean ± SD) of indicated protoplasts after 10-min hypotonic shock of indicated conditions. n, cell number; *P*-values, two-tailed t-test. (F) Working model of Ca^2+^-dependent PM remodeling events controlling massive hypoosmotic PM expansion.

In brief, the activation of MscS-like channel-mediated Ca^2+^ influx and increased water permeability was assumed to occur above a threshold PM strain (ε), set to 0.01. ^47^ Rather than explicitly simulating random protoplast rupture events, we highlighted strain regions above a threshold of ε = 0.06, ^48^ where the likelihood of rupture increases with higher ε values. We also introduced an osmoregulation module to depict the flux of osmolytes by modifying a previous model, ^47^ and a similar Ca^2+^ homeostasis module to simulate Ca^2+^ levels by MscS-like channels and intracellular calcium pumps. Based on many other studies, ^9,34,49-51^ we presumed that increased PM tension (σ, a linear function of ε) accelerated exocytosis (k_1_) and eisosome disassembly (k_3_) while attenuated endocytosis (k_2_) and eisosome reassembly (k_4_). Besides, high Ca^2+^ levels likely enhanced exocytosis albeit with an unclear mechanism, as the PM constantly expanded more with Ca^2+^ in buffers than without it under modest hypoosmotic shock (ΔC = − 0.8 M sorbitol) in *tcb1Δtcb3Δ* protoplasts, some of which managed to survive even without external Ca^2+^ (Figure S4A). Thus, the exocytic rate k_1_ was simulated to depend on both σ and Ca^2+^ levels with a maximum rate of 8 μm^2^/min (discussed below). Tcb1/3-mediated lipid transfer (k_5_) in *WT* was estimated at a constant rate of 1.5 μm^2^/min upon Ca^2+^ influx, equivalent to moving 3.5 PL molecules per second per Tcb1-Tcb3 complex, which is within the estimated rate range for *in vivo* lipid transport. ^52^ We then monitored hypotonic PM expansion across various scenarios involving *WT*, related mutants or drug treatment by alternating associated parameters.

Our model reproduced *WT* protoplast expansion during acute hypoosmotic shock, featuring the rapid PM surface increase accompanying Ca^2+^ influx and the variation in PM strain until osmotic homeostasis (Figure 4B). It also showed that exocytosis was the major contributor to surface increment among various membrane sources during expansion (Figure S4B). Consistent with experiment, the model exhibits successive Ca^2+^ influx peaks in surviving cells (Figures 4B and S4C). More importantly, simulated protoplasts enter a vulnerable period when ε surpassed 0.06, with rupture likelihood increasing at higher ε values (framed in Figure 4B). Such simulated vulnerable periods occurred earlier and were longer, exhibiting much higher ε in simulations lacking Tcb1/3-mediated lipid transfer, exocytosis, or Ca^2+^ influx (Figures 4C and S4D). This indicates an increased risk of quick cell rupture with limited PM expansion under these conditions, highly resembling our experimental data (Figures 1A, 1B, 1D, S1A, 2A-C, 2E, S2C, 3B, 3D, and S3E). The model also reproduced the delay in Ca^2+^ influx (when ε exceeds 0.01) without endocytosis, mimicking LatA treatment, as the lack of endocytosis slows PM tension increase (Figures 4C and S4D). The eisosome contribution in the model was adjusted to be small enough such that it behaves like *WT* (Figures 4C and S4D). Remarkably, our model recapitulated a lower risk of protoplast rupture with Tcb1/3 overexpression due to enhanced strain reduction, without affecting predictions of other scenarios, when introducing a time delay (t_d_) between the activation of Tcb1/3-mediated lipid transfer and Ca^2+^-dependent acceleration of exocytosis (Figures 4C and 4D). Otherwise, Tcb1/3 overexpression would mostly survive the acute hypoosmotic shock, inconsistent with our observations (Figure 3F). This delay is also supported by our evidence that *S. pombe* likely lacks Ca^2+^ sensors for directly boosting vesicle trafficking and fusion. Likewise, with this delay, our model effectively predicted the trends in survival and expansion for both *WT* and *tcb1Δtcb3Δ* protoplasts under moderate hypotonic shocks (i.e., ΔC = −0.9 M or −0.8 M sorbitol; Figures 4D, 4E and S4E). This also suggests that a short delay of ∼5 seconds would likely mirror physiological conditions, at least with our parameters (Figure 4D). Simulations also highlighted the importance of the rapid activation of Tcb1/3-mediated lipid transfer upon Ca^2+^ influx for relieving membrane strain critical for PM integrity. This role seems crucial for cells to survive other PM-damaging conditions, such as heat stress (Figure S4F). In summary, our model successfully recapitulated protoplast expansion under all tested conditions during hypotonic shock, suggesting that PM tension- and Ca^2+^-dependent regulation of hitherto recognized key remodeling events may represent the main mechanism underlying hypoosmotic PM expansion for osmotic balance (Figure 4A).

Altogether, we revealed following PM remodeling events during hypotonic shock: 1) the stretch-activating MscS-like channels mediate Ca^2+^ influx upon initial expansion driven by hypoosmotic pressure; 2) Ca^2+^-dependent E-Syts-mediated lipid transfer rapidly supplies PLs to ensure the integrity of the expanding PM; 3) increased exocytosis, stimulated by both high PM tension and elevated Ca^2+^, delivers bulk membranes to support rapid and massive PM expansion for osmotic equilibrium (Figure 4F). Importantly, our numerical simulations suggest that a combination of rapid non-vesicular lipid transport and bulk exocytic membrane inflow establishes a robust mechanism for keeping PM integrity during dramatic cell surface/volume alteration.

## Discussion

Physiological adaption to rapid cell expansion passively resulting from environmental stimuli, such as acute hypoosmotic shock, is a challenging process requiring prompt and coordinated cellular responses to maintain homeostasis and prevent cell integrity loss. Here, we postulate that Ca^2+^-triggered non-vesicular lipid transport is necessary as a swift remedy for fission yeast to ensure PM integrity before the arrival of other machineries required for expansion or repair under severe stresses. This E-Syts-mediated role might be an undeveloped strategy before the emergence of animal synaptotagmins in fungi and plants, where mechanical support from cell walls largely avoids drastic cell surface and volume expansion. Similar to our observations, malfunction of plant E-Syts (albeit with confusable names as synaptotagmins) also engenders cell lysis under abiotic and biotic stresses. ^53^ Somewhat in line with this idea, Ca^2+^-dependent dramatic hypotonic expansion in HeLa cells did not require E-Syts (data not shown), where synaptotagmins might plausibly stimulate rapid exocytosis. It is also an intriguing coincidence that yeast and plant cells have relatively abundant ER-PM contacts facilitating such direct lipid transfer. ^16,18,28,54^

Unlike *in vitro* studies where both homo- or heterodimers of E-Syts seem active in lipid transfer, ^22,23,43^ *S. pombe* Tcb1 and Tcb3 with comparable endogenous levels appear to function mainly as hetero-complexes *in vivo*. Tcb1-Tcb3 association stabilizes their accumulation at ER-PM contacts, essential for effective lipid transfer. A previous study suggests that tricalbins make highly curved cER, narrowing ER-PM spacing to 7-8 nm and promoting lipid transfer. ^25^ Thus, the rapid cER reticulation^34^ and Ca^2+^ influx seen during hypotonic stock should result in highly robust lipid transfer. We believe that E-Syts-mediated role identified here is mainly to maintain PM integrity rather than to increase PM surface area with following evidence: 1) extensive PM expansion was also seen in cells with compromised activity or lower levels of E-Syts (Figures 3F and S3D) and 2) some *tcb1Δtcb3Δ* protoplasts showed *WT-*like massive PM expansion during moderate hypotonic shocks (Figures 4E and S4A). Although we think that membrane supply from such direct lipid transfer accounts for some small PM surface increase, half of PLs delivered to the cytoplasmic PM leaflet must be flopped to the outer leaflet to make a proper PM bilayer. Hence, floppases and/or scramblases are anticipated to aid the process. Unlike TMEM16 scramblase, its yeast homolog Ist2 that lacks lipid scrambling activity^55^ is dispensable for hypotonic PM expansion (data not shown). It remains to be seen if floppases are involved. Alternatively, spontaneous incorporation of PLs into both PM leaflets exposed at small wounds might occur with physical forces (e.g., tension).

Despite our results suggest a Ca^2+^-dependent exocytosis enhancement in *S. pombe* protoplasts, it does not completely contradict the long-held view in budding yeast that exocytosis is constitutive and Ca^2+^-independent. ^56-58^ Consistent with these early data testing Ca^2+^ necessity for post-Golgi vesicle fusion, we do not think that this Ca^2+^-mediated regulation works on vesicles at late secretory stages; it might involve vesicle generation. As coated vesicle formation is believed to occur in the order of seconds, ^59-61^ our estimated ∼5 s delay in Ca^2+^-related exocytic acceleration might be reasonable. Such links between Ca^2+^ and exocytosis in yeast still await elucidation. Moreover, our quantitative analysis suggests that the accelerated exocytosis rate significantly exceeds its constitutive speed in walled cells (∼1.24 μm^2^ /min equivalent to fusion of ∼ 40 vesicles per min, estimated from *S. pombe* growth rate). We reason that PM sites for exocytosis in protoplasts are no longer limited to tips in walled cells and increasing ER-free PM zones become available for exocytic fusion as the cortical ER unfolds into a reticulated meshwork during hypotonic expansion. ^34^ Thus, the basal exocytosis capacity in protoplasts is expected to increase and we estimated it to be maximally ∼6 to 7 times of that in walled cells, ∼8 μm^2^/min. High Ca^2+^ levels might instead facilitate early secretory pathways to indirectly support such increased exocytosis. A typical fission yeast protoplast contains around 900 secretory vesicles, ^62^ equal to ∼28 μm^2^, which may ideally serve as an adequate membrane pool for bulky PM expansion. During our manuscript preparation, a similar study in budding yeast showed that massive hypoosmotic PM expansion was ATP-independent, which may have led authors to overlook exocytic membrane delivery. ^63^ In contrast, hypotonic PM expansion of *S. pombe* protoplasts was highly sensitive to ATP shortage (Figure S4G).

Notably, opposite to what was observed in protoplasts, hypoosmotic cell expansion of walled fission yeast is more pronounced in *msy1Δmsy2Δ* than in *WT* with higher cytosolic Ca^2+^ levels in the mutant. ^13^ Although different channels are responsible for hypotonic Ca^2+^ influx in walled cells, the mechanism of Ca^2+^-regulated expansion is likely the same. Unlike spheroplasts that require Ca^2+^ influx to passively survive hypoosmotic expansion, drastic cell surface and volume increase restricted by rigid walls may become detrimental in walled cells, necessitating tight control of cytosolic Ca^2+^ levels. Excessive Ca^2+^ is also toxic and can cause apoptosis-like yeast cell death. ^64^ Homeostatic regulation of Ca^2+^ in this regard is also vital for protoplasts to survive hypotonic shock, thus cytoplasmic Ca^2+^ levels always returned to baseline in survivors. Different from MscS-like channels, the other stretch-activated channels shown in Figure S1C contain large extracellular domains which may require cell wall for their activities. This may partly explain the prevalent role of MscS-like channels in mediating Ca^2+^ influx in protoplasts. Similar to these proposed functions, MSL10, the MscS-like homolog in *Arabidopsis thaliana*, senses PM tension and controls cytosolic Ca^2+^ spikes in response to hypotonic cell swelling. ^14^

In conclusion, our work elucidates the mechanism how Ca^2+^-dependent vesicular and non-vesicular lipid transfer ensures PM integrity and its expansion to survive hypoosmotic cell swelling. As key players in Ca^2+^-dependent PM remodeling events are highly conserved in fungi and plants, similar mechanism may widely apply to other cell types.

## Acknowledgments

Many thanks are due to Q. Chen, S. Oliferenko and M. Balasubramanian for sharing strains used in this work. B.M. and D.Z. were funded by the Temasek Life Sciences Laboratory. D.R. and D.V. were funded by NIH grant R35GM136372.

## Author Contributions

B.M. and D.Z. designed and performed experiments. D.R. and D.V. designed and performed simulations. G.G. optimized the microfluidics setup. B.M., D.R., D.Z. and D.V. analysed the data. B.M. and D.Z. wrote the first draft; D.R. and D.V. wrote the mathematical model section; all of them contributed to the manuscript.

## Materials and Methods

### Strain construction

All fission yeast strains used in this study are listed in Supplementary Table 1. Strains were constructed from laboratory stocks using standard methods. The phosphatidylinositol-4-phosphate (PI_4_P) marker PH_Osh2_-GFP^16^, the phosphatidylinositol-4,5-biphosphate (PI_4,5_P_2_) marker PH_Num1_-GFP^16^ and the phosphatidic acid (PA) marker PABD_Spo20_-GFP were constitutively expressed from *ura4* locus under *rtn1* promoter. The phosphatidylserine (PS) marker GFP-Lact-C2^65^ was expressed from *ade6* locus under *act1* promoter, the sterol marker GFP-D4H and mCherry-D4H^66^ were expressed from *leu1* locus under *pil1* promoter, and the diacylglycerol (DAG) marker C1β-GFP^67^ was expressed from *ura4* locus under *cdc15* promoter. Cytosolic Ca^2+^ reporter^27^ was expressed from *leu1* locus under *adh1* promoter. C-terminally GFP, mCherry, nYFP, or cYFP tagged Msy1, Msy2, Tcb1, Tcb2, Tcb3 and Pil1 were expressed from respective native locus. *tcb3*^*WWW*^ was made by substituting all 251L, 417Y and 419L with tryptophan, and *tcb1*^*WW*^ was made by substituting 300I and 468Y with tryptophan. The truncation of each functional domain of Tcb1 or Tcb3 at respective native loci was made by deleting corresponding amino acids except for *tcb1ΔC2D*, where the entire C-terminus from 1333V was truncated. Overexpression of tricalbins were made by switching the native promoter with the thiamine repressible promoter *Pnmt1* at respective native loci.

### Media and growth conditions

For all experiments unless otherwise stated, *S. pombe* cells were grown exponentially at 24 °C in YES (yeast extract with supplements) or EMMS (Edinburgh minimal media supplemented with appropriate amino acids) before specific treatments or imaging. Gene expression under *nmt1* promoter was induced in EMMS for at least 20 hours (h). EMMS with 5 mg/mL thiamine (Thi) (Sigma-Aldrich, T1270-25G) was used to suppress the expression. Latrunculin A (LatA; ThermoFisher, L12370) dissolved in dimethyl sulfoxide (DMSO) was used at a final concentration of 50 μM. Brefeldin A (Sigma-Aldrich, B6542-5MG) dissolved in ethanol was used at a final concentration of 100 μg/mL.

#### Protoplast preparation

*S. pombe* cells were grown in YES for about 18 h. 1 OD_595_ of cells were harvested and washed three times with EMMS without CaCl_2_ (EMMS-Ca^2+^). Cells were digested at 24 °C for 1 h in EMMS-Ca^2+^ supplemented with 5 mg/mL lysing enzyme (Sigma-Aldrich, L3768-1G), 3 mg/mL zymolase (nacalai tesque, Zymolyase® −20T) and 1.2 M D-sorbitol (hereafter named as the steady-state isotonic buffer) to generate protoplasts. Protoplasts were kept in the steady-state isotonic buffer for 30 minutes to ensure their osmotic equilibrium before hypoosmotic treatment. For experiments related to tricalbins overexpression, cells were firstly grown in EMMS for over 20 h and then in YES culture for 1 h before enzymatic removal of cell walls.

#### Drug and heat stress treatment

For drug treatment experiments, BFA or LatA was added to spheroplasts solution, treatment buffer, and steady-state media to a final concentration of 50 μM or 100 μg/mL respectively. Before loaded to the chamber for the hypoosmotic shock, spheroplasts were pretreated with LatA for 10 minutes (min) or with BFA for a specified time (i.e., 0, 10, 20, 30, 40, 50, 60, 70, 80, or 90 min). For ATP depletion treatment, 20 mM 2-deoxyglucose (2-DG) and 10 μM antimycin A, or 5 mM NaN_3_ and 5 mM NaF dissolved in EMMS without glucose was added to treatment buffer, and steady-state media and spheroplasts solution just (less than 5 mins) before hypoosmotic shock. For heat stress experiment, 2 OD_595_ of cells from fresh culture were collected and concentrated in 100 μL YES, then incubated at 42°C for 2 h before adding Propidium Iodide (PI) to a final concentration of 20 μg/mL. After 5 min incubation with PI at 42°C, cells were imaged at 24 °C.

### Microscopy

#### Microfluidic based spinning disk confocal microscopy

Spheroplasts loading and solution exchange were performed with OB1 pressure pump (Elveflow). The microfluidics chamber (CellASIC ONIX plate, Y04E-01-5PK) was perfused with the steady-state isotonic buffer at a pressure of 1 psi for 1h to build a homeostatic environment. Spheroplasts were loaded to the chamber with 4.5 μm height at 2.6 psi. Imaging started at the time when the steady-state isotonic buffer was switched to respective treatment buffer (i.e., media supplemented with 0.2/0.3/0.4 M D-sorbitol with 5/1/0.5/0.1 mM of CaCl_2_) at 1.16 psi, capturing a single focal plane at cell center or periphery (for Pil1-mCherry in Fig. S2A) every 3 seconds for 10 min. After each experiment, the solution in the chamber was exchanged with the steady-state buffer to rebuild the homeostasis.

Scanning confocal microscopy for Figures S2B, S3A, and S3C was performed on an Olympus FLUOVIEW FV3000 (Olympus Corporation, Japan) equipped with a U Plan Super Apochromat 100X, 1.4 NA Oil immersion objective lens, a 488 nm (for GFP excitation) and a 561 nm (for mCherry excitation) solid state laser, with high sensitivity-spectral detectors. Typically, we acquired image stacks that consisted of 9 z-sections with 0.5 μm spacing (55.2 nm per pixel). Walled cell imaging was performed on *S. pombe* cells placed in sealed growth chambers containing 2% agarose YES medium.

Cells shown in Figures 1A, 1D, S1A, 2A, 2D, 2E, S2A, S2F, 3C, S3B, S3C, S3E, S4C, and S4F were imaged using Nikon TiE system (CFI Plan Apochromat VC 100X, 1.4 N.A. objective) equipped with Yokogawa CSU-X1-A1 spinning disk unit, Prime 95B™ Scientific CMOS camera (110 nm per pixel) and a DPSS 491 nm 100 mW, 514 nm 100 mW and DPSS 561 nm 50 mW laser illumination under the control of MetaMorph Premier Ver. 7.7.5. Spheroplasts shown in Figures 1E and 3C were imaged in the 4.5μm height microfluidic chamber with the Live SR module (Gataca systems).

### Protein level estimation and Co-immunoprecipitation (Co-IP)

For protein level estimation in Fig. S3D, cells were grown to log phase, harvested with freshly prepared IP buffer (50 mM Tris-HCl, 150 mM NaCl, 2 mM EDTA, 50 mM NaF, 0.1 mM Na_3_VO_4_, 1 mM PMSF, 1.5 mM Benzamidine-HCl, and Roche protease inhibitor cocktail) 1% Nonidet P (NP)-40. Cells were mixed with glass beads and homogenized in a Mini Bead Beater (Biospec, OK, USA) at 4 °C and total cell lysates were harvested after removing cell debris. Cell lysates were then subjected western blotting probed by mouse anti-RFP (6g6, Chromotek, IL, USA) and mouse anti-α-tubulin antibodies (TAT-1, a gift from K. Gull, University of Oxford, UK). For detection, IRDye800 conjugated anti-mouse was used, and subsequent quantification analyses were done on the Odyssey infrared imaging system (LI-COR, NE, USA) and ImageJ 1.48v software package (NIH, MD, USA).

For co-immunoprecipitation in Fig. 3D, glass bead-homogenized cells were suspended in 300 μL of IP buffer with 1% Nonidet P (NP)-40, at 4°C and total cell lysates were harvested after removing cell debris. Typically, lysates were adjusted to the same total protein concentration, diluted with IP buffer and precleared for non-specific binding by incubation with Dynabeads Protein G (Invitrogen, CA, USA) for 1 h at 4°C. The pre-cleared lysate was then subjected to incubation with rabbit anti-GFP (Abcam, MA, USA) for 1 h at 4°C and then with Dynabeads Protein G for 45 min at 4°C. Beads were washed five times with IP buffer. Samples were resuspended in SDS-loading buffer and subjected SDS-PAGE and western blotting probed by rabbit anti-GFP, mouse anti-RFP antibodies as above. For detection, IRDye800 conjugated anti-mouse and IRDye680 conjugated anti-rabbit antibodies were used.

### Image Analysis

Calculation of relative surface area, max *rCa*^2+^_*cyto*_, Pearson’s R, and time difference (Δt) Cell area at the medial focal plane for each time point was traced and measured using ImageJ “Analyze Particles” function. The surface area (S) and volume (V) of the protoplast at each time point were calculated as *S* = 2*π*^2^*r*(*R − r*) + 4*πr*^2^ + 2*π*(*R − r*)^2^ and 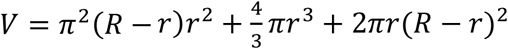, respectively, where R is the radius of the medial focal plane, and r is half of the chamber height (2.25 μm). Relative surface area was calculated as 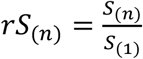.

The expansion profile along time was plotted using MATLAB with moving average (n=5). The cytosolic Ca^2+^ level 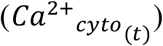was determined by analyzing the mean grey value of the medial plane at each time point (t). The relative total cytosolic calcium level (*rCa*^2+^_c*yto*_) at a given time point (t) was calculated as 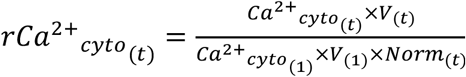

Here, *rCa*^2+^_*cyto*(*t*)_represents the relative total calcium content, not the concentration, in the specific protoplast volume. *Ca*^2+^_*cyto*_ and *V*_(1)_denote the grey value and volume at the first point of the 10-min movie, respectively, while 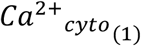 and *V*_(*t*)_refer to the grey value and volume at the t^th^ time point. *Norm*_(*t*)_, the normalization factor at the t^th^ time point, was introduced for photobleaching correction, and was obtained by normalizing *Ca*^2+^_*cyto*_ thatwere averaged from ten *WT* protoplasts under the same imaging settings without hypoosmotic treatment.

For each analyzed cell, apparent volumetric flux was defined as 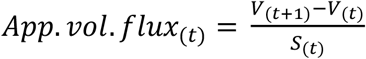 and 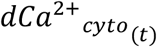 was set as 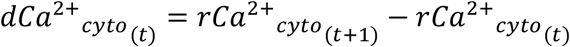. The Pearson’ R of the cell was computed by MATLAB function corrcoef() using *App. vol*. f*lux* and *dCa*^2+^_*cyto*_ datasets during the first calcium elevation period, which is from the time point one to the time point before increased *rCa*^2+^_*cyto*_ firstly dropped.

Dt_WT_ and Δt_LatA_ shown in Fig. S2D were defined as the duration between the start of surface area increment and the time point with visible increase in *rCa*^2+^_*cyto*_. In our MATLAB algorithms, the start of surface area increment was defined as the first time point where its and the next 9 consecutive values of *dS*_(*t*)_= *S*_(*t*+1)_− *S*_(*t*)_were positive, and the threshold for visible *rCa*^2+^_*cyto*_ increase was set as 2, which was determined by averaging such *rCa*^2+^_*cyto*_ values from 30 visually scored cells.

#### Kymograph generation

Kymographs were generated using ‘MultipleKymograph’ plugin in ImageJ with the indicated regions of 10-pixel width.

### Description of computational model

Definition and values of all variables used in the model are listed in Supplementary Table 2.

#### Protoplast model shape

We assume the protoplast is compressed within the microfluidic chamber of height *h* = 4.5 μm, in contact with the top and bottom chamber surfaces. The contact areas are assumed to be circular regions of radius *R*_*m*_, connected to each other by plasma membrane in the shape of the outer half of a torus with major radius *R*_*m*_ and minor radius *r* = h/2. Here, *R*_*m*_ = *R* − r where *R* is the radius of the medial focal plane as defined in the main text. The area of this pancake-like shape can be written as 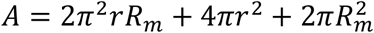while the volume as 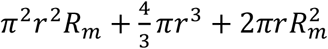. During protoplast expansion, we evolve *R*_m_ through time, assuming the protoplast remains cylindrically symmetric. Initially *R*_*m*_ = 0.79 μm, corresponding to a protoplast with surface area 103 μm^2^ similar to the approximate average starting area of the protoplast in experiments.

#### Protoplast expansion

The volume of the protoplast, *V*, changes through time due to water flux across the plasma membrane^3,47,68^ and growth:

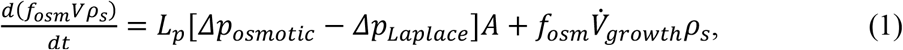

where *f*_*osm*_ is the fraction of cell volume that is osmotically active (described below), *A* is the surface area of the protoplast, the first term on the right-hand side describes the flow of water across the membrane due to the imbalance between osmotic and hydrostatic pressure while the second describes cell growth with volumetric rate 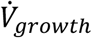The hydraulic conductivity, *L*_*p*_, can also be written as *L*_*p*_ = *Pρ*_*s*_*v*_*s*_/*ℛT* where *P* is the membrane permeability, *ρ*_*s*_ is the mass density of water, *ℛ* is the ideal gas constant, and *T* is the temperature. ^3,68^ Using *Pv*_*s*_ as a parameter instead of *L*_*p*_, the value of *ρ*_*s*_ does not appear in the final equations of the model.

Some differences of our model of protoplast expansion to prior models of osmotic regulation of yeast protoplast cell volume are: (1) we study the kinetics of protoplast expansion while prior models focused on the equilibrium between osmotic and hydrostatic pressure, (2) we don’t account for changes in nuclear volume, and (3) we consider large hypotonic perturbations, which bring the protoplast far from the ideal osmometer regime. ^69,70^ The changes of cell volume under the 1.2 M to 0.2 M external concentration changes are several-fold smaller than what would be expected from ideal osmometer behavior, indicating that leakage of osmolytes has to be taken into consideration for our experiments (see below). Our model reproduces ideal osmometer response after equilibration over a few minutes, for osmotic perturbations less than 0.3 M and provided the water permeability is sufficiently high to allow volume equilibration within that time interval (it is possible that the experiments in previous studies^69,70^, which were started in lower external sorbitol concentration, were close to the high water permeability value, *P*_*high*_, in our model, explained below, which would be sufficient to see such a response).

#### Osmotic pressure difference

Using the Van’t Hoff equation, the osmotic pressure difference can be written as^3,47^

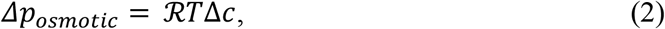

where Δ*c* is the difference in osmolyte concentration between the inside and outside of the protoplast (for simplicity we only consider a single concentration of osmolyte^47^, which we consider to include both ionic and non-ionic components). Reduction of the external osmolarity leads to an increase in Δ*c* and an increase in cell volume from Equation 1.

#### Hydrostatic pressure

Hydrostatic Laplace pressure is the difference in fluid pressure between the inside and outside of a closed, curved surface that develops due to plasma membrane tension. The Laplace pressure on the torus region can be written in terms of the two radii of curvature^31,47^

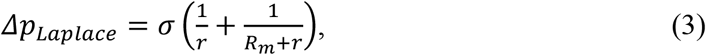

where *r* and *R*_*m*_ + r are the two principal radii of curvature and σ is the membrane tension. The pressure on the flat region of the protoplast shape is supported mechanically by the walls of the microfluidic chamber. Additionally, we assume the tension is equilibrated throughout the entire membrane surface and so both the torus and flat regions have the same σ. The membrane tension is related to the plasma membrane surface area as

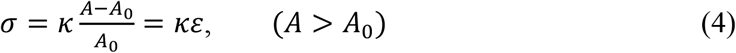

where *κ* is the effective area extension modulus of the plasma membrane, *ε* is the membrane strain, and *A*_0_ is the zero-strain area of the protoplast. In this model we use *κ* = 0.09^3^ N/*m*, which represents a combined modulus of both unfolding of membrane and direct elastic stretching of the membrane^3^ (see reason for this exact value in Initial conditions below).

#### Osmotically active volume fraction

We assume the fraction of the cell volume that is osmotically active is *f*_*osm*_ = 0.7. The remaining osmotically inactive fraction represents the “dry volume” of the cell and includes proteins and other solid material within the cell. ^69,71,72^ The left-hand side of Equation 1 is the change in the mass of water within the protoplast. The dry volume fraction within the cell remains at 1 − f_*osm*_ by processes not explicitly modeled.

#### Protoplast growth

We assume there is a constant addition of volume to the cell over time that accounts for the background growth of the cell and that this growth is, for simplicity, independent of the osmolyte concentration gradient across the cell. When the growing shape is spherical:

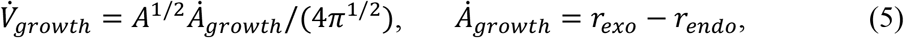

where 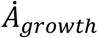 is the corresponding area growth rate and *r*_*exo*_, r_*endo*_ are the wild-type exocytosis and endocytosis rates of membrane delivery for a growing cell tip of walled *S. pombe*. ^73^ This growth rate approximately matches our experimentally observed rates of protoplast growth.

#### Initial conditions

The initial concentration of osmolytes inside the protoplast is assumed to be at a steady state value slightly larger than the external 1.2 M concentration (i.e. the protoplast osmolarity is assumed to have largely equilibrated with the external concentration before the hypoosmotic shock). The initial tension is set to σ_*ss*_ = 5 × 10^−4^ N/m to match measurements of protoplast tension at steady state^37^ (at 0.8 M external concentration). The initial osmolyte concentration difference that balances the Laplace pressure caused by *σ*_*ss*_ is Δ*c*_*ss*_ = 0.15^4^ mol/m^3^. Our results are similar if both *σ*_*ss*_ and Δ*c*_*ss*_ are zero instead. The effective area expansion modulus *κ* was set to a value such that the initial membrane strain of protoplasts at steady state has a relatively low value *ε*_*SS*_ = 0.005^4^. The initial value of *A*_0_(*t* = 0)is fixed by *κ*, the initial area, *A*(*t* = 0), and the initial value of the membrane tension, *σ*_*ss*_, through Equation 4.

#### Membrane area transfer between reservoirs

Except for small expansions, the volumetric growth of the protoplast upon hypoosmotic stress must be accompanied by membrane area delivery or otherwise the cell will burst^1^. We model the transfer of membrane to *A*_0_ (the undeformed zero strain area of the protoplast plasma membrane) from three internal reservoirs of membrane area (Fig. 4A). In addition to keeping the membrane tension low, this membrane transfer serves to attenuate the increase in σ and therefore reduces the Laplace pressure in Equation 1. The internal membrane reservoirs are membrane that can be delivered via exocytosis (*A*_*ves*_), eisosomes (*A*_*eiso*_), and tricalbin-associated membrane reservoir (*A*_*tcb*_). These reservoirs of membrane can transfer to and from *A*_0_, depending on the area membrane strain, *ε*, and the level of excess Ca^2+^, Δ*Ca*, namely the Ca^2+^ concentration above the steady state cytoplasmic concentration (which is assumed to be much smaller than the external Ca^2+^ concentration). These reservoirs do not include growth terms, and we assume the inclusion of such terms would be a small, second-order correction to the model. The area balance equation for *A*_0_ includes all the membrane transfer rates included in the model and is given as

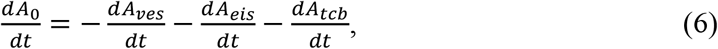

with each of these terms described in more detail below.

Exocytosis and endocytosis

The area of the exocytic vesicle reservoir, *A*_*ves*_, changes due to both exocytosis (*k*_1_)and endocytosis (*k*_2_)as

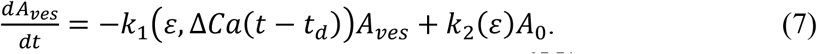

The exocytosis rate *k*_1_ is assumed to increase with membrane strain^37,74^ linearly, to leading order approximation. We further assume that the response to increase in strain has Ca^2+^-dependent and Ca^2+^-independent components:

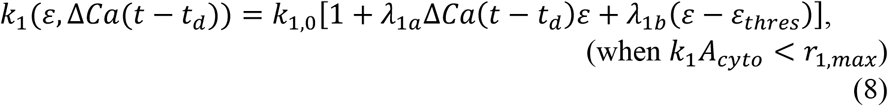

where *t*_*d*_ is a time delay in the response of exocytosis to the internal Ca^2+^ signal above the steady state value, and *λ*_1*a*_, *λ*_1*b*_, *ε*_*thres*_ are constants. We note that the dependence of the second term on the right-hand side on both the Ca^2+^ concentration and the membrane strain ensures the exocytosis rate does not drive the protoplast to negative strains (which occurs if *A* < A_0_). We also assume the Ca^2+^-independent pathway increases exocytosis above the threshold strain *ε*_*thres*_ of increased permeability (discussed below). If *k*_1_*A*_*cyto*_ > r_1,*max*_ by the above equation, then we set *k*_1_*A*_*cyto*_ = r_1,*max*_ = 8 μm^2^/m*in* be an upper limit on the rate of exocytosis (see main text).

The strain dependence for the endocytosis rate, *k*_2_, is

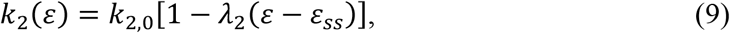

where *k*_2,0_ is the reference rate at steady-state value of strain, *ε*_*ss*_, and *λ*_2_ is a constant describing the increase or decrease of endocytosis with strain above or below *ε*_*ss*_. ^37,74,75^

The steady state value, *A*_*cyto*_(*t* = 0), is set to 30% of the initial surface area of the protoplast as justified in the Discussion section in the main text. The steady state rate of exocytosis is *k*_1,0_*A*_*cyto*_(*t* = 0)= r_*exo*_ and of endocytosis is *k*_2,0_*A*_0_(*t* = 0)= r_*endo*_. For a concentration drop of 1 M, the resulting increase in exocytosis is about six times its steady state value while the endocytosis rate drops near zero. These two changes to exocytosis/endocytosis with strain are of similar magnitude to extrapolated values of prior experiments at smaller concentration drops. ^37^

#### Eisosome disassembly and reformation

The amount of area within the eisosome reservoir changes due to both eisosome disassembly and reformation as

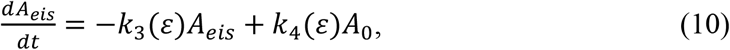

where both rates are dependent on membrane strain. ^5,37,51^ The strain dependence of the eisosome disassembly rate is

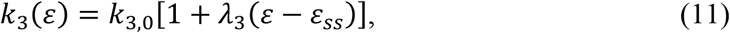

where *k*^3^,_0_ is the reference rate at steady-state value of strain and *λ*^3^ is a constant. Similarly, the strain dependence of the eisosome reformation rate is

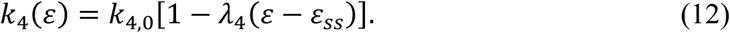

Both *k*^3^ and *k*^4^ scale with *ε* − *ε*_*ss*_ because we assume that the eisosome deformation and reformation process is directly influenced by any deviation from the steady state membrane strain. ^5,37,51^ Initially *A*_*eis*_(*t* = 0)is set to be 5% of the surface area of the protoplast. ^5,37^

#### Tricalbin membrane delivery

The pool of area for tricalbin associated delivery (*A*_*tcb*_) is assumed to be far from being exhausted because the ER is the store for this membrane delivery, which is assumed to be much larger than the other two reservoirs. Tricalbin delivery is assumed to be triggered by internal Ca^2+^ in a stepwise manner as

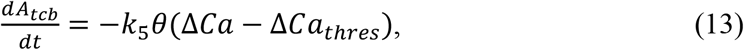

where *k*_5_ is the rate of membrane delivery by tricalbins, *θ* is the Heaviside step function and Δ*Ca*_*thres*_ is a Ca^2+^ activation threshold.

#### Strain dependence of permeability

The early stages of experimental protoplast growth due to hypoosmotic stress have two phases: an initial, slow growth phase of around 1-2 minutes followed by a much faster growth phase (Fig. 1A). Assuming the entire concentration gradient is experienced by the cell at the beginning of the slow expansion, the only variable in Equation 1 that allows for this large change in water influx is the permeability, *P*. Consequently, we assume the water permeability constant, *P*, in our model has two values: one for low strain (*P*_*low*_) and one for high strain (*P*_*high*_), with the threshold strain that triggers the switch between these two values being *ε*_*thres*_ = 0.01 as written below.

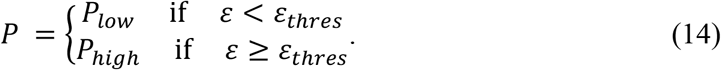

We hypothesize that this increase of the plasma membrane permeability could occur due to opening of mechanosensitive pores, Ca^2+^ channels, or other nanopores within the plasma membrane upon significant membrane stretching.

#### Leakage of osmolytes

The model also includes a flux of osmolytes (ionic and non-ionic components other than Ca^2+^, which is described below) across the membrane through either mechanosensitive channels or active transport. ^75^ These two osmolyte fluxes modulate the internal osmolyte concentration, *c*_*in*_, as

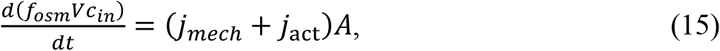

where *c*_*in*_ = Δ*c* + c_*out*_, *j*_*mech*_ is the passive flux through mechanosensitive channels, and *j*_act_ is the flux through active transport. We neglect the increase of *c*_*in*_ due to growth, which we assume to be a second-order correction to our model. The mechanosensitive channels open in response to stress and allow for the passive flux of ions through the channels in the direction of the concentration gradient. The passive ion flux can be written as^47^

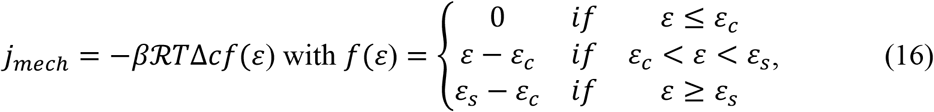

where *β* is a rate constant, and *f*(*ε*) is a piecewise function dependent on strain, where *ε*_*c*_ is a threshold strain above which mechanosensitive channel begin opening and *ε*_*s*_ is the saturation strain above which mechanosensitive channels remain open. The active flux represents pumping of ions that return the cell to its initial, steady state osmolyte concentration difference at long times. Specifically,

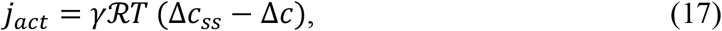

where *γ* is a constant controlling the rate of strain independent flux through active pumps. *j*_*act*_ is largely negligible for the value of *γ* chosen and on the timescale of the simulation (∼10 min), consistent with slow recovery over an hour of fission yeast protoplasts experiencing hypertonic perturbations. ^69^

#### Ca^2+^ influx

We assume the Ca^2+^ level within the cell is controlled by both a passive and active flux in the same manner as the osmolyte concentration as

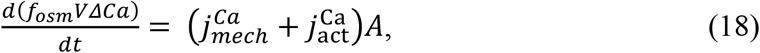

where *ΔCa* is the difference in the internal and external level of Ca^2+^. We do not seek a precise model of Ca^2+^ concentration in the cell, which involves additional complex dynamics beyond the scope of our model. Since the concentration of Ca^2+^ within the protoplast is small compared to *c*_*in*_, we neglect the contribution of Ca^2+^ to the osmotic pressure. We further note that the reported units of Ca^2+^ can be rescaled by adjusting the model parameters that depend on Δ*Ca*: *λ*_1*a*_ in Equation 8 and the threshold of the Heaviside term in *k*_5_ of Equation 13, which were chosen empirically to match experimental observations.

The mechanosensitive channels for Ca^2+^ are assumed to be all-or-nothing and so depend on the strain with a step function as

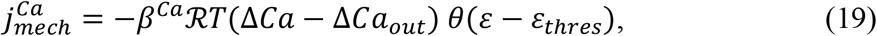

where Δ*Ca*_*out*_ is the difference in Ca2+ concentration outside the cell and the inside of the cell at steady state and *β*^*Ca*^ is a rate constant controlling the strain independent Ca^2+^ flux through mechanosensitive channels. The active Ca^2+^ pumps also attempt to return the concentration difference of Ca^2+^ across the membrane to its steady state value as

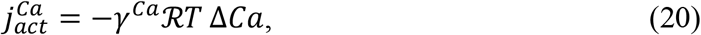

where *γ*^*Ca*^ is a constant controlling the rate of strain independent Ca^2+^ flux through active transport. Unlike with the osmolyte active pumping,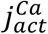 is large on the timescale of the simulation and accounts for the rate of decrease in the Ca^2+^ level after the plateau in Fig. 4B, which is roughly on a similar timescale as that in experiment Fig. S4C. Here, we write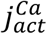 as representing flux of Ca^2+^ out of the cell, though pumping of Ca^2+^ from the cytoplasm into the ER would generate the same effect.

#### Membrane rupture

If the area strain is higher than *ε*_*rupture*_ = 0.06, the model cell is assumed to be in danger of bursting. ^48,76^ Regions where the model protoplast experiences strains above this rupture threshold are indicated by dashed lines in Figures 4B, 4C, 4D, S4D, and S4E.

#### Differential equation solver

We numerically solve the equivalent form of Equation 1 as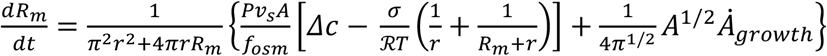which derives from previous equations in addition to the coupled equations describing membrane delivery (Equations 6, 7, 10, 13) and osmolyte/Ca^2+^ leakage (Equations 15, 18). We assume Δ*Ca* is zero at all *t*<0. We used a simple explicit Euler scheme as implemented in Matlab to solve our system of differential equations with a timestep of 0.01 s over a timeframe of 600 s. The Matlab code will be made available after publication.

#### Assumptions with mutants and drug treatments

We investigate several protoplast expansion scenarios related to mutants or drug treatments with our model (Fig. 4C); we describe the assumptions in the model for each case here. For *pil1Δ* case, we set *A*_*eiso*_(*t* = 0)to zero. For *tcb1/3Δ* case, we set *k*_5,*max*_ to zero. For BFA treatment, we set both the exocytosis and endocytosis rates to zero (i.e. *k*_1,0_ = k_2,0_ = 0). For no Ca^2+^ influx case, we set the external Ca^2+^ level, Δ*Ca*_*out*_, to zero. For *tcb1/3* overexpression (OE) case, we double *k*_5,*max*_ to be 3 μm^2^/min. For LatA treatment, we set endocytosis rate to zero (*k*_2,0_ = 0). The rate of exocytosis is also reduced in the LatA treatment case, as previously suggested. ^37,77^ The exocytosis rate was specifically reduced to half, *k*_1,0_ = r_1,*max*_/2*A*_*cyto*_(*t* = 0), by matching the predicted delay of reaching the threshold strain to experiment (Fig. S2D). Without a reduction, this delay becomes longer than 10 minutes and no longer matches experiment because without endocytosis the value of *A*_0_(*t*)increases faster with time than in *WT*.

### Statistical evaluation

Types of statistical tests, sample sizes, definition of error bars and measured values were provided either in figures or figure legends. Typically, student’s t test was used to generate *P*-values which were computed using Microsoft excel.

## Code availability

Bastian Bechtold (2016), Violin plots for Matlab, Github project, https://github.com/bastibe/Violinplot-Matlab.

**Supplementary Table 1.**
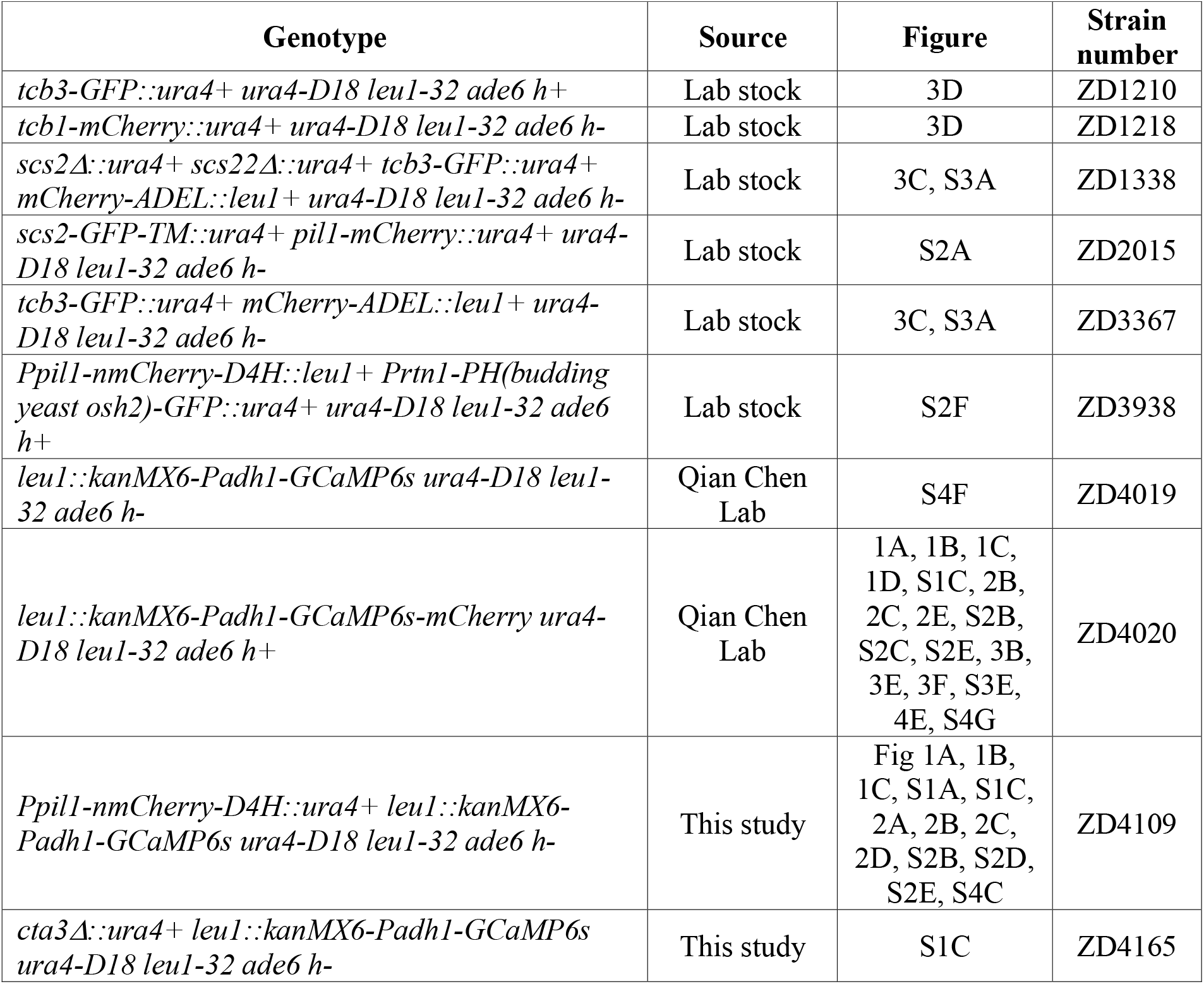
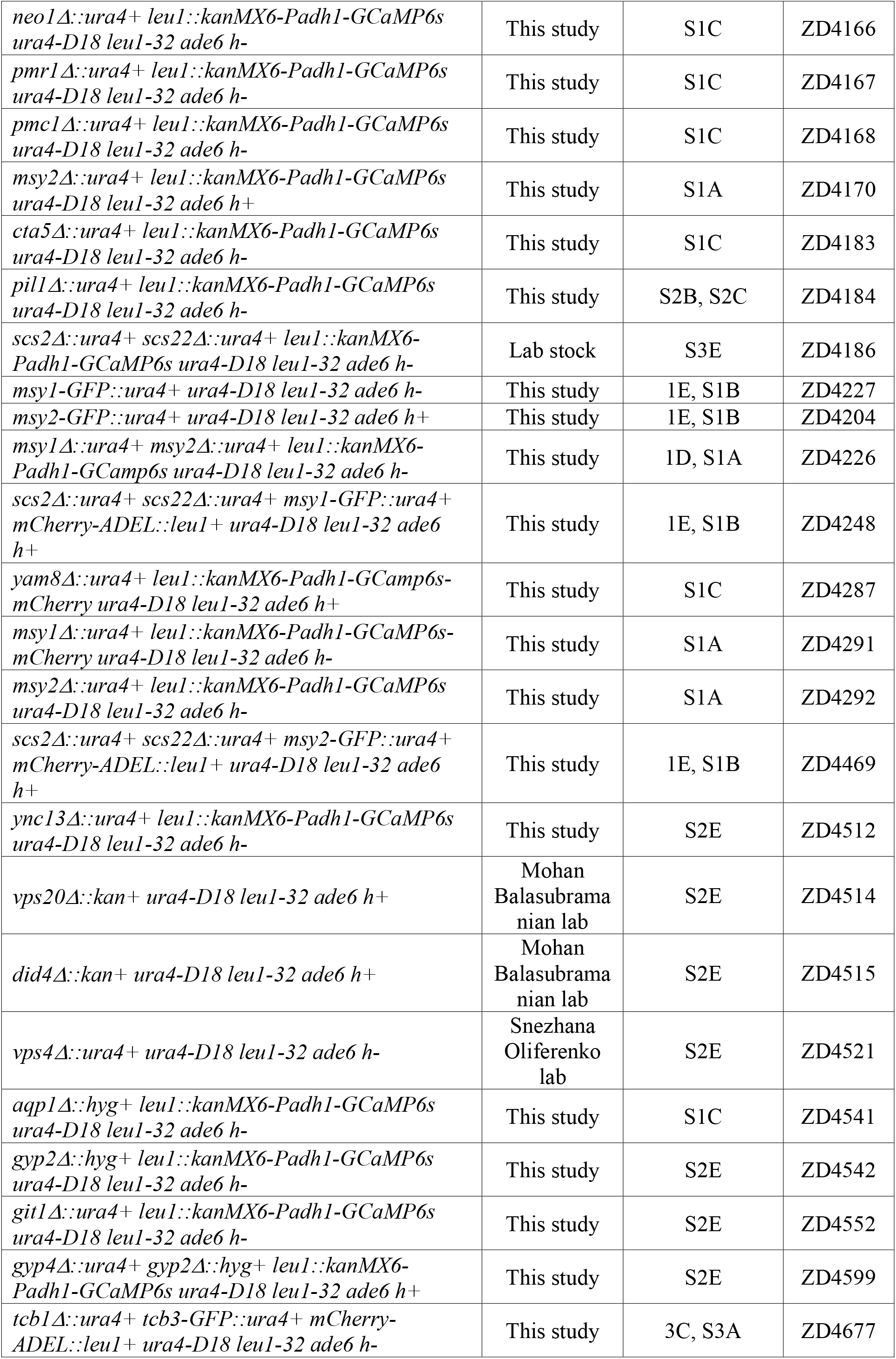
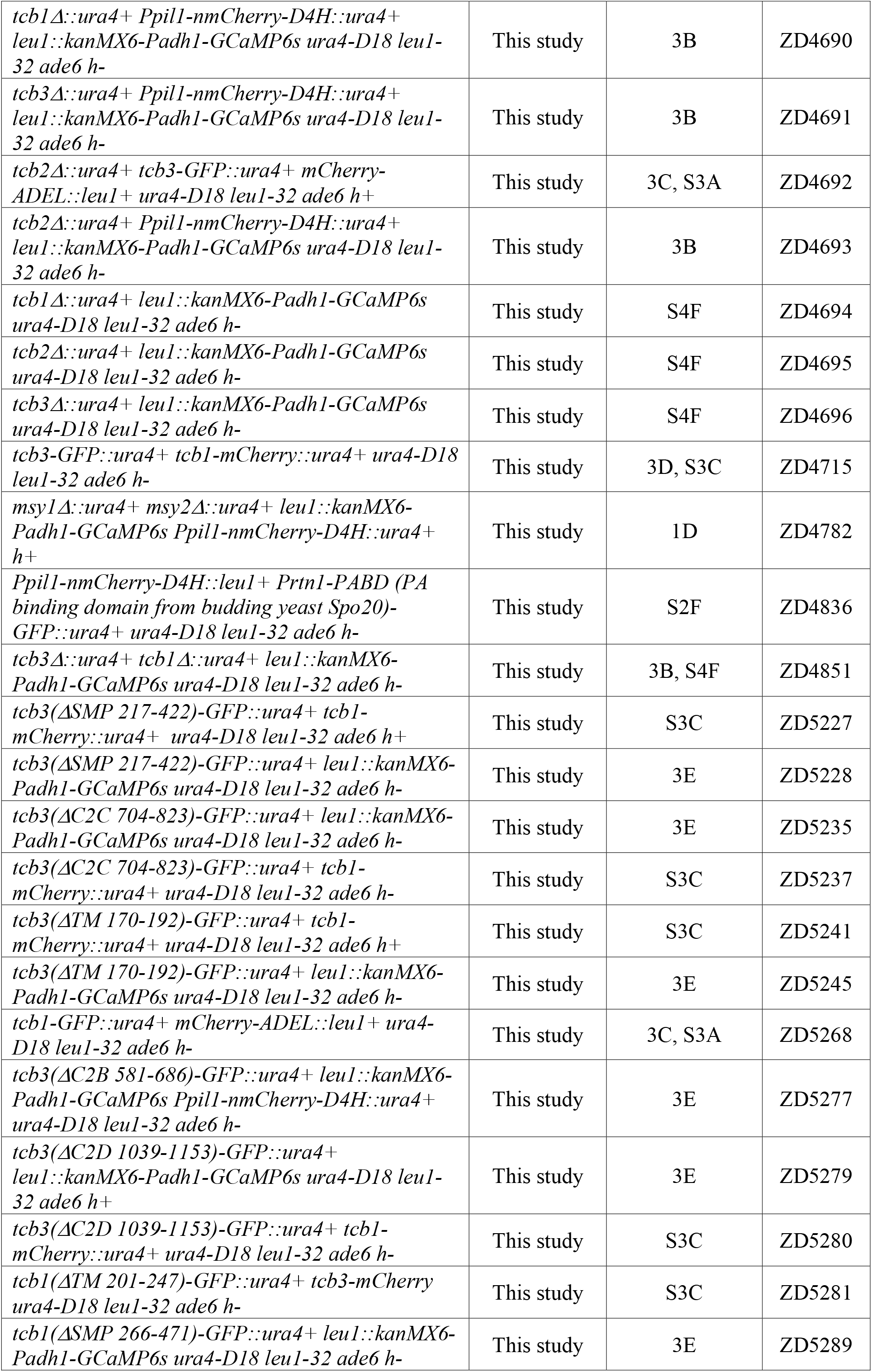
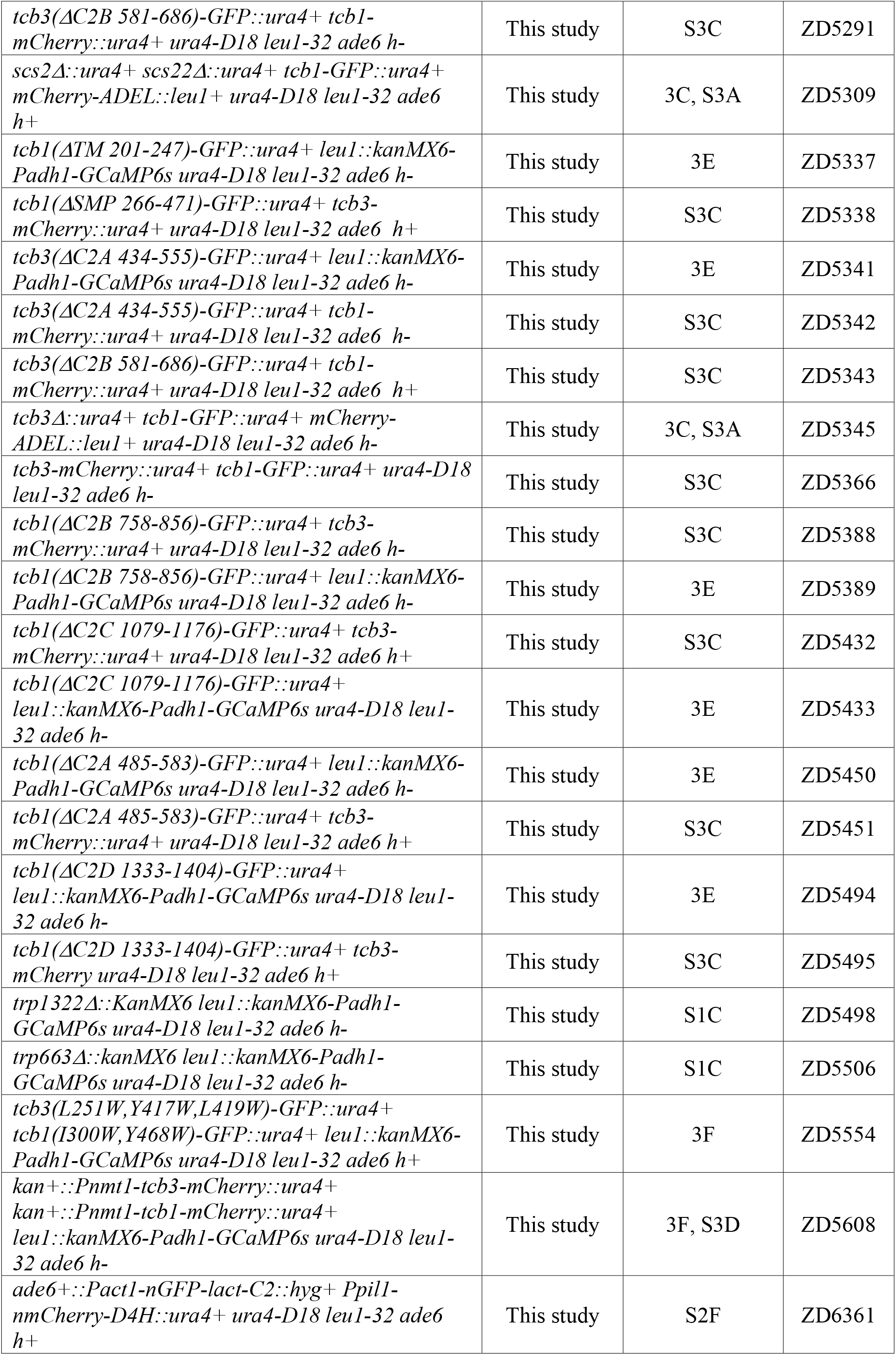
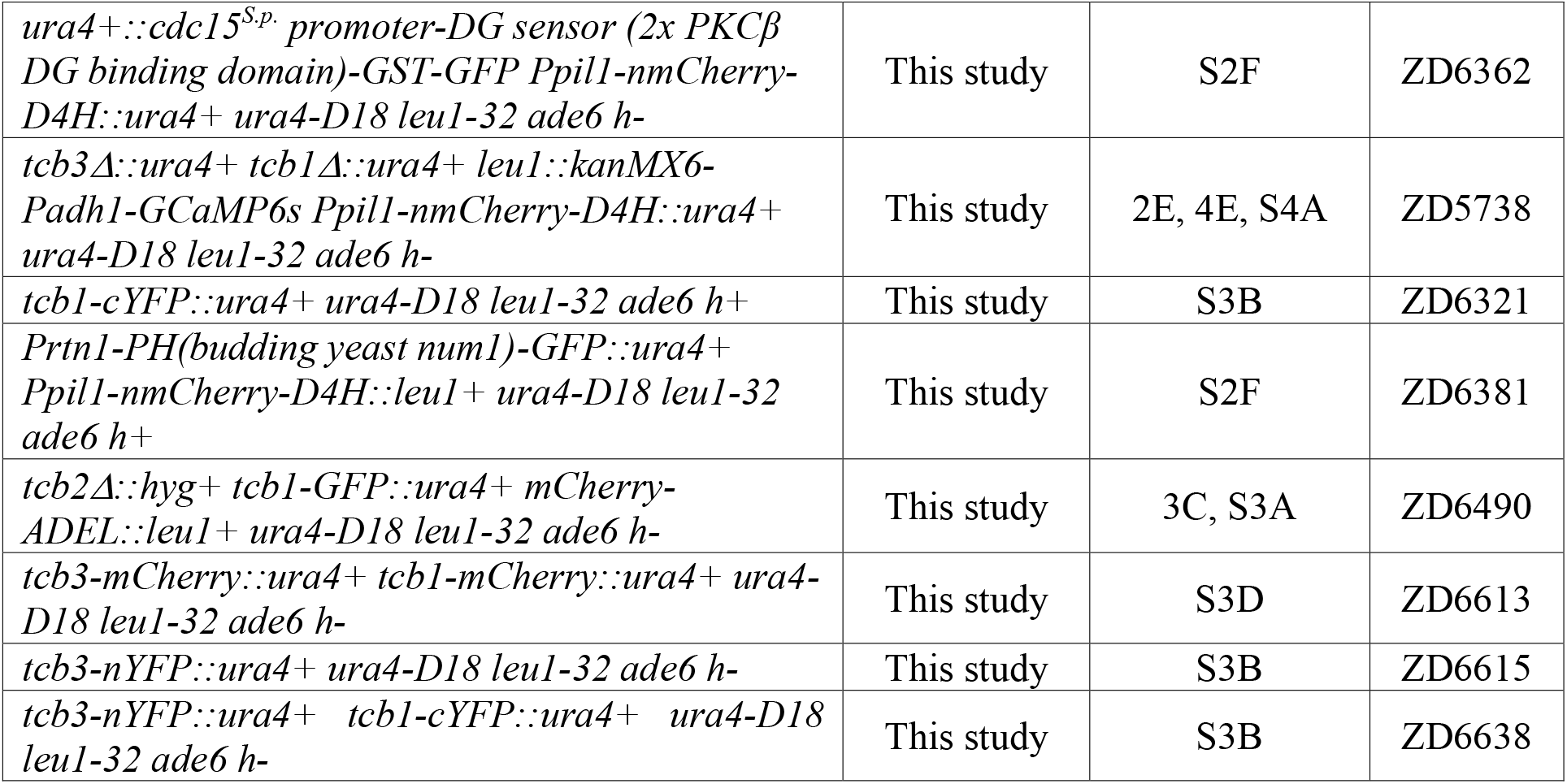

**Supplementary Table 2.**

| Variable | Value | Description |
| --- | --- | --- |
| $h$ | 4.5 $\mu\text{m}$ | Height of microfluidic chamber |
| $r$ | 2.25 $\mu\text{m}$ | Minor radius of torus in confined protoplast shape |
| $R_m(t = 0)$ | 0.79 $\mu\text{m}$ | Initial radius of contact circle with microfluidic walls; major radius of torus |
| $\kappa$ | 0.093 N/m | Stretching modulus of protoplast membrane; set to give a low initial membrane strain ($\varepsilon = 0.0054$) |
| $f_{osm}$ | 0.7 | Osmotically active fraction of the protoplast volume |
| $A_0(t = 0)$ | 102 $\mu\text{m}^2$ | Initial area of the zero-strain protoplast; fixed by $A(t = 0)$ and $\sigma(t = 0)$ |
| $A_{ves}(t = 0)$ | 30.8 $\mu\text{m}^2$ | Initial area within the cytoplasmic reservoir of exocytic vesicles primed for delivery |
| $A_{eiso}(t = 0)$ | 5.14 $\mu\text{m}^2$ | Initial area in eisosome reservoir; set to 5% of $A(t = 0)$ |
| $\sigma_{ss}$ | $5 \times 10^{-4}$ N/m | Initial membrane tension of protoplast<sup>37</sup> |
| $\Delta c_{ss}$ | $1.54 \times 10^{-1}$ mol/m<sup>3</sup> | Determined from $\sigma_{ss}$ and Equations 2 and 3 |
| $\Delta Ca_{out}$ | 2.5 mol/m<sup>3</sup> | Initial difference between outside and inside Ca<sup>2+</sup> level |
| $\Delta Ca_{thres}$ | 1.0 mol/m<sup>3</sup> | Threshold level of Ca<sup>2+</sup> for tricalbin membrane delivery, estimated |
| $P_{low}$ | $4.88 \times 10^{-9}$ m/s | Low strain water permeability of plasma membrane; varied to fit initial slope of protoplast expansion |
| $P_{high}$ | $2.31 \times 10^{-7}$ m/s | High strain water permeability of plasma membrane; close to value reported for *S. cerevisiae*,<sup>78</sup> but smaller than values reported for bare lipid membranes;<sup>3,79</sup> varied to fit initial slope of protoplast expansion |
| $\varepsilon_{thres}$ | 0.01 | Threshold membrane strain for switching from $P_{low}$ to $P_{high}$ |
| $\varepsilon_c$ | 0.01 | Threshold membrane strain for opening of mechanosensitive ion channels, 1/5 of that used in the previous study<sup>47</sup> to allow for channels to open significantly before reaching rupture threshold of 0.06 |
| $\varepsilon_s$ | 0.03 | Saturation membrane strain for mechanosensitive ion channels, triple that of $\varepsilon_c$ as in the previous study<sup>47</sup> |
| $\varepsilon_{Ca}$ | 0.01 | Threshold membrane strain for $Ca^{2+}$ influx into protoplast |
| $r_{exo}$ | $1.284 \mu m^2/min$ | Single tip exocytosis rate<sup>73</sup> |
| $r_{endo}$ | $0.942 \mu m^2/min$ | Single tip endocytosis rate taken<sup>73</sup> |
| $r_{1,max}$ | $8 \mu m^2/min$ | Maximum rate of exocytosis, estimated |
| $k_{1,0}$ | $6.94 \times 10^{-4} 1/s$ | Zero strain rate of exocytosis; set to give $r_{exo}$ for the initial exocytosis rate |
| $k_{2,0}$ | $1.54 \times 10^{-4} 1/s$ | Zero strain rate of endocytosis; set to give $r_{endo}$ for the initial endocytosis rate |
| $k_{3,0}$ | $1.62 \times 10^{-4} 1/s$ | Zero strain rate of eisosome disassembly; set in conjunction with $k_{4,0}$ to observe some eisosome removal with slow reformation on 10 min timescale |
| $k_{4,0}$ | $8.15 \times 10^{-6} 1/s$ | Zero strain rate of eisosome reformation; set in conjunction with $k_{3,0}$ to observe some eisosome removal with slow reformation on 10 min timescale |
| $k_{5,max}$ | $1.5 \mu m^2/min$ | Rate of tricalbin membrane delivery when strain exceeds $\varepsilon_{thres}$ |
| $\lambda_{1a}$ | $750 m^3/mol$ | Constant controlling increase of exocytosis rate with internal $Ca^{2+}$ and strain, estimated |
| $\lambda_{1b}$ | 100 | Constant controlling increase of exocytosis rate with strain alone, estimated |
| $\lambda_2$ | 15 | Constant controlling decrease of endocytosis rate with strain, estimated |
| $\lambda_3$ | 60 | Constant controlling increase of eisosome disassembly with strain, estimated |
| $\lambda_4$ | 15 | Constant controlling decrease of eisosome reformation with strain, estimated |
| $t_d$ | 5 s | Time delay of exocytosis rate response to internal $Ca^{2+}$ signal |
| $\beta$ | $3.1 \times 10^{-10} mol m^{-2} Pa^{-1} s^{-1}$ | Constant controlling rate of osmolyte influx through passive mechanosensitive channels; gives 10x the flux used in the previous study<sup>47</sup> for a $10^4$ larger concentration difference, gives approximate amount of osmolyte leakage to match to experimental expansion |
| $\gamma$ | $1.4 \times 10^{-15} mol m^{-2} Pa^{-1} s^{-1}$ | Constant controlling rate of osmolyte influx/efflux through active ion channels; larger than the value used in the previous study<sup>47</sup> but small enough that it does not affect our simulation results significantly |
| $\beta^{Ca}$ | $7 \times 10^{-11} mol m^{-2} Pa^{-1} s^{-1}$ | Constant controlling rate of $Ca^{2+}$ influx through passive mechanosensitive channels; approximated to give $Ca^{2+}$ spike behavior |
| $\gamma^{Ca}$ | $7 \times 10^{-12} mol m^{-2} Pa^{-1} s^{-1}$ | Constant controlling rate of $Ca^{2+}$ influx/efflux through active ion channels; approximated to give $Ca^{2+}$ spike behavior |
| $\Delta t$ | 0.01 s | Timestep for Euler solver |

