## Supplementary Material for "Ca^2+^-dependent vesicular and non-vesicular lipid transfer controls hypoosmotic plasma membrane expansion"

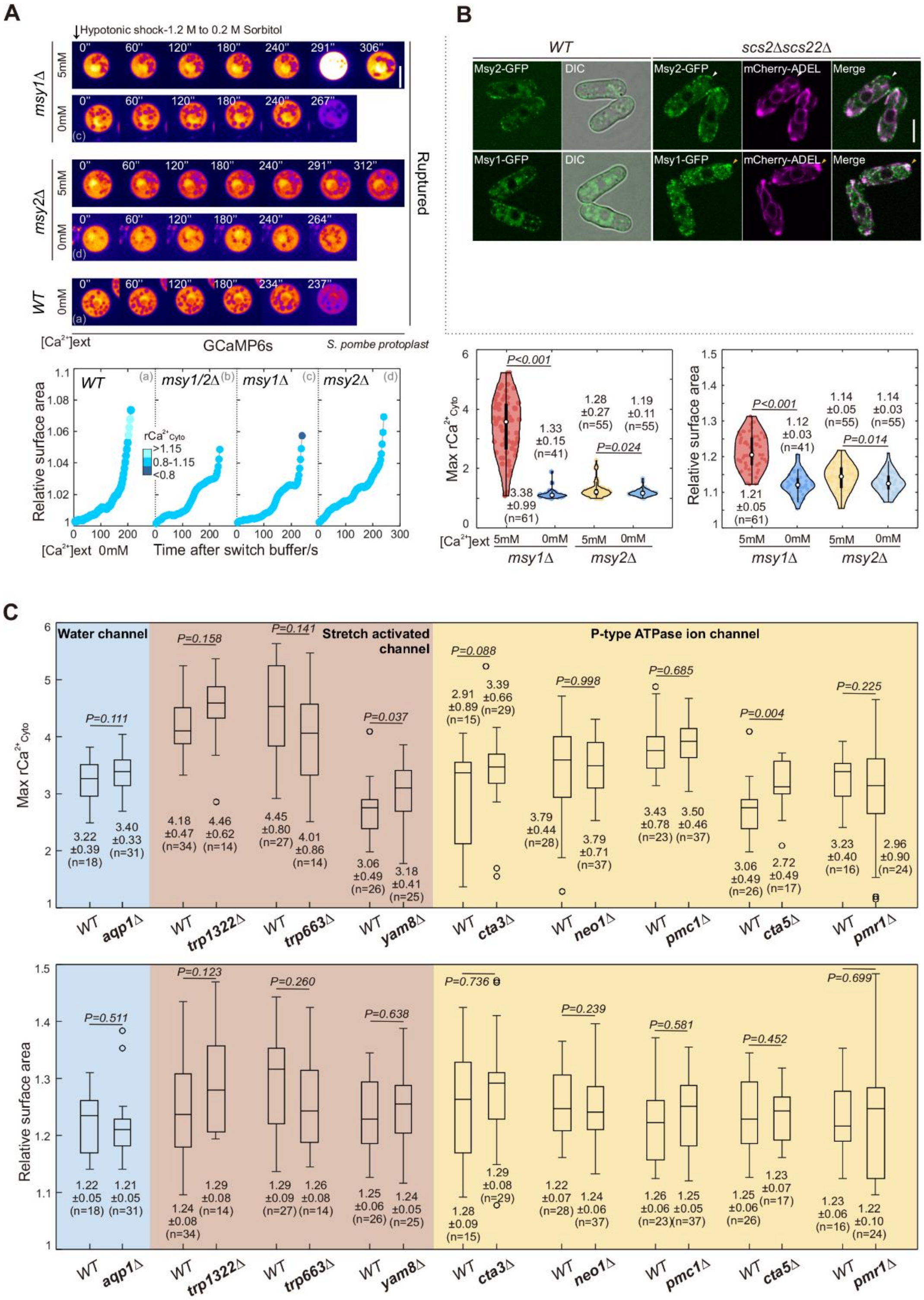
A
Hypotonic shock-1.2 M to 0.2 M Sorbitol
msy1Δ
5mM
0mM
msy2Δ
WT
Ruptured
[Ca2+]ext
GCaMP6s
S. pombe protoplast
Relative surface area
WT
msy1/2Δ
msy1Δ
msy2Δ
rCa2+Cyto
>1.15
0.8-1.15
<0.8
[Ca2+]ext 0mM
Time after switch buffer/s
B
WT
scs2Δscs22Δ
Msy2-GFP
DIC
mCherry-ADEL
Merge
Msy1-GFP
Max rCa2+Cyto
P<0.001
P=0.024
3.38 ±0.99 (n=61)
1.33 ±0.15 (n=41)
1.28 ±0.27 (n=55)
1.19 ±0.11 (n=55)
P<0.001
P=0.014
1.21 ±0.05 (n=61)
1.12 ±0.03 (n=41)
1.14 ±0.05 (n=55)
1.14 ±0.03 (n=55)
[Ca2+]ext 5mM 0mM 5mM 0mM
msy1Δ
msy2Δ
C
Water channel
Stretch activated channel
P-type ATPase ion channel
Max rCa2+Cyto
P=0.111
P=0.158
P=0.141
P=0.037
P=0.088
P=0.998
P=0.685
P=0.004
P=0.225
3.22 ±0.39 (n=18)
3.40 ±0.33 (n=31)
4.18 ±0.47 (n=34)
4.46 ±0.62 (n=14)
4.45 ±0.80 (n=27)
4.01 ±0.86 (n=14)
3.06 ±0.49 (n=26)
3.18 ±0.41 (n=25)
2.91 ±0.89 (n=15)
3.39 ±0.66 (n=29)
3.79 ±0.44 (n=28)
3.79 ±0.71 (n=37)
3.43 ±0.78 (n=23)
3.50 ±0.46 (n=37)
3.06 ±0.49 (n=26)
2.72 ±0.49 (n=17)
3.23 ±0.40 (n=16)
2.96 ±0.90 (n=24)
WT aqp1Δ
WT trp1322Δ
WT trp663Δ
WT yam8Δ
WT cta3Δ
WT neo1Δ
WT pmc1Δ
WT cta5Δ
WT pmr1Δ
Relative surface area
P=0.511
P=0.123
P=0.260
P=0.638
P=0.736
P=0.239
P=0.581
P=0.452
P=0.699
1.22 ±0.05 (n=18)
1.21 ±0.05 (n=31)
1.24 ±0.08 (n=34)
1.29 ±0.08 (n=14)
1.29 ±0.09 (n=27)
1.26 ±0.08 (n=14)
1.25 ±0.06 (n=26)
1.24 ±0.05 (n=25)
1.28 ±0.09 (n=15)
1.29 ±0.08 (n=29)
1.22 ±0.07 (n=28)
1.24 ±0.06 (n=37)
1.26 ±0.06 (n=23)
1.25 ±0.05 (n=37)
1.25 ±0.06 (n=26)
1.23 ±0.07 (n=17)
1.23 ±0.06 (n=16)
1.22 ±0.10 (n=24)
WT aqp1Δ
WT trp1322Δ
WT trp663Δ
WT yam8Δ
WT cta3Δ
WT neo1Δ
WT pmc1Δ
WT cta5Δ
WT pmr1Δ

**Figure S1 MscS-like channels control the extracellular $Ca^{2+}$ influx during hypotonic PM expansion.** (A) Time-lapse spinning disk confocal images of representative protoplasts expressing GCaMP6s with indicated hypotonic shocks. Shown are pseudo-colored images at central focal planes. Times, relative to buffer switch. Graph shows relative PM surface expansion curves of (a)-(d) protoplasts with colors denoting total relative cytosolic $Ca^{2+}$ level (r$Ca^{2+}$Cyto). (b) is shown in Figure 1D. Quantifications of maximum r$Ca^{2+}$Cyto and maximum relative surface area (mean ± standard deviation [SD]) of indicated protoplasts after 10-min acute hypotonic shock of indicated conditions are also included. (B) Central focal plane spinning disk confocal images of indicated cells. Arrows, GFP signals in ER-free PM regions. Scale bar, 5 µm. (C) Box-and-whisker plot of maximum r$Ca^{2+}$Cyto and maximum relative surface area (mean ± standard deviation [SD]) of indicated protoplasts after 10-min acute hypotonic shock of indicated conditions. n, cell number; *P*-values, two-tailed t-test.

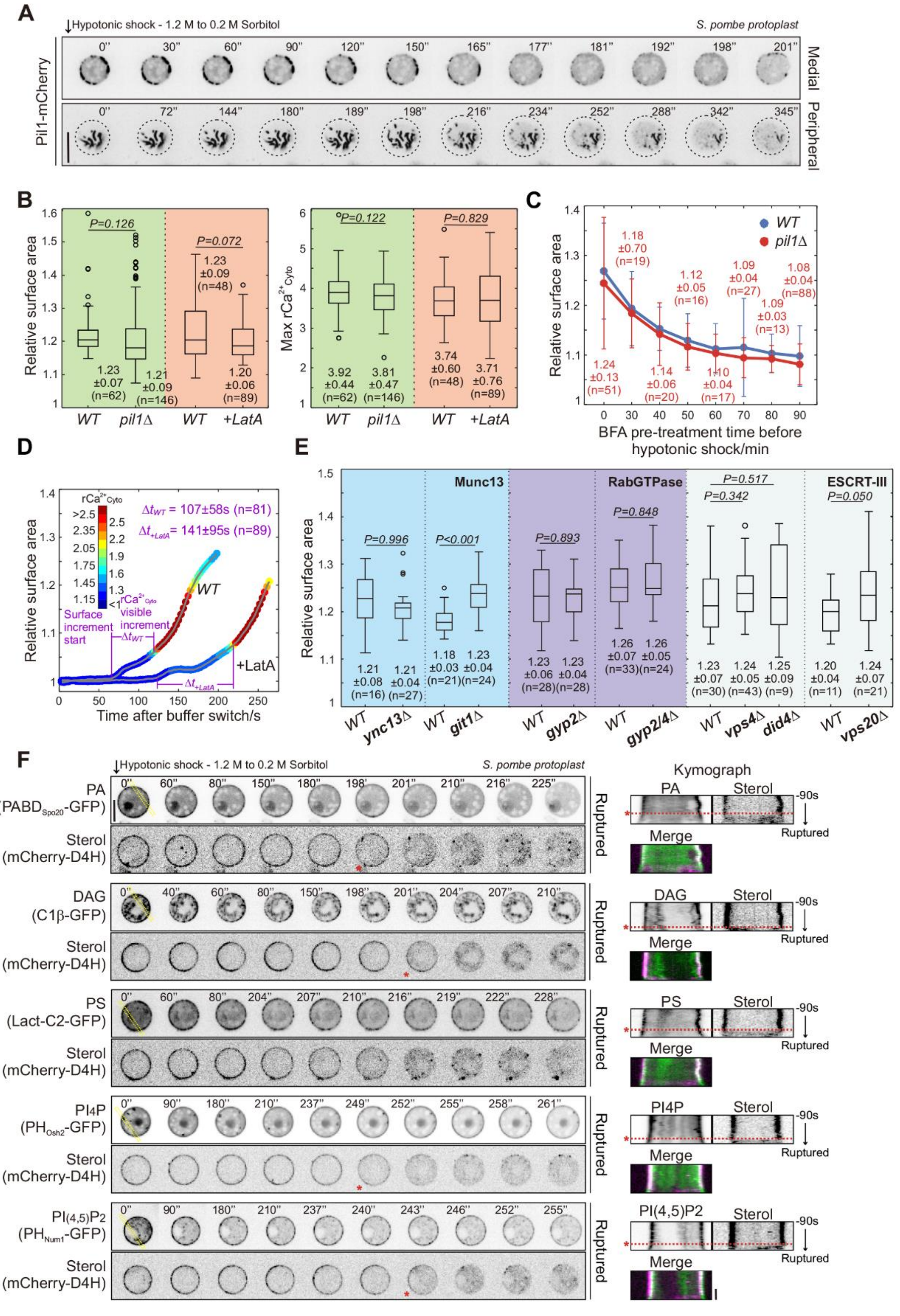

A
Hypotonic shock - 1.2 M to 0.2 M Sorbitol
S. pombe protoplast
Pil1-mCherry
Medial
Peripheral
B
Relative surface area
P=0.126
P=0.072
WT pil1Δ WT +LatA
Max rCa2+Cyto
P=0.122
P=0.829
C
WT
pil1Δ
BFA pre-treatment time before hypotonic shock/min
D
Relative surface area
Time after buffer switch/s
Δt_WT = 107±58s (n=81)
Δt_+LatA = 141±95s (n=89)
Surface increment start
rCa2+Cyto visible increment
+LatA
E
Munc13
RabGTPase
ESCRT-III
P=0.996
P<0.001
P=0.893
P=0.848
P=0.517
P=0.342
P=0.050
WT ync13Δ WT git1Δ WT gyp2Δ WT gyp2/4Δ WT vps4Δ did4Δ WT vps20Δ
F
Hypotonic shock - 1.2 M to 0.2 M Sorbitol
S. pombe protoplast
PA (PABDSpo20-GFP)
DAG (C1β-GFP)
PS (Lact-C2-GFP)
PI4P (PHOsh2-GFP)
PI(4,5)P2 (PHNum1-GFP)
Sterol (mCherry-D4H)
Ruptured
Kymograph
Merge
-90s

**Figure S2 Eisosomes disassembly and endocytosis play minor roles in massive hypotonic PM expansion.**

(A) Time-lapse spinning disk confocal images at central and peripheral focal planes of representative protoplasts expressing Pil1-mCherry with indicated hypotonic shocks. Dash lines indicate the cell outline. Times, relative to buffer switch. (B and E) Box-and-whisker plot of maximum relative surface area and/or maximum r$Ca^{2+}$Cyto (mean ± SD) of indicated protoplasts after 10-min acute hypotonic shock of indicated conditions. *P*-values, two-tailed t-test. (C) Quantifications of maximum relative surface area (mean ± SD shown for *pil1Δ*) of BFA-pretreated *WT* and *pil1Δ* protoplasts after 10-min acute hypotonic shock. The relative surface areas of *WT* from time point 0 min to 90 min are 1.27±1.0 [n=34], 1.19±0.7 [n=10], 1.15±0.4 [n=15], 1.13±0.5 [n=16], 1.11±0.5 [n=17], 1.12±1.0 [n=26], 1.10±0.4 [n=12], and 1.10±0.6 [n=19] (mean ± SD). Error bars, 2 × SD. (D) The relative PM surface expansion curves of corresponding protoplasts with colors denoting r$Ca^{2+}$Cyto. Δt indicates time difference (mean ± SD) from start of surface expansion to the visualization of apparent $Ca^{2+}$ influx. n, cell number. (F) Time-lapse spinning disk confocal images at central focal planes of representative protoplasts expressing indicated lipid probes together with mCherry-D4H under indicated hypotonic shocks. Times, relative to buffer switch. Kymographs were generated from 90 seconds before the protoplast rupture. Yellow boxes, the regions selected for the kymograph. Red asterisks and dashed lines, the time point for initiation of sterol retrograde movement. Scale bar, 5 µm.

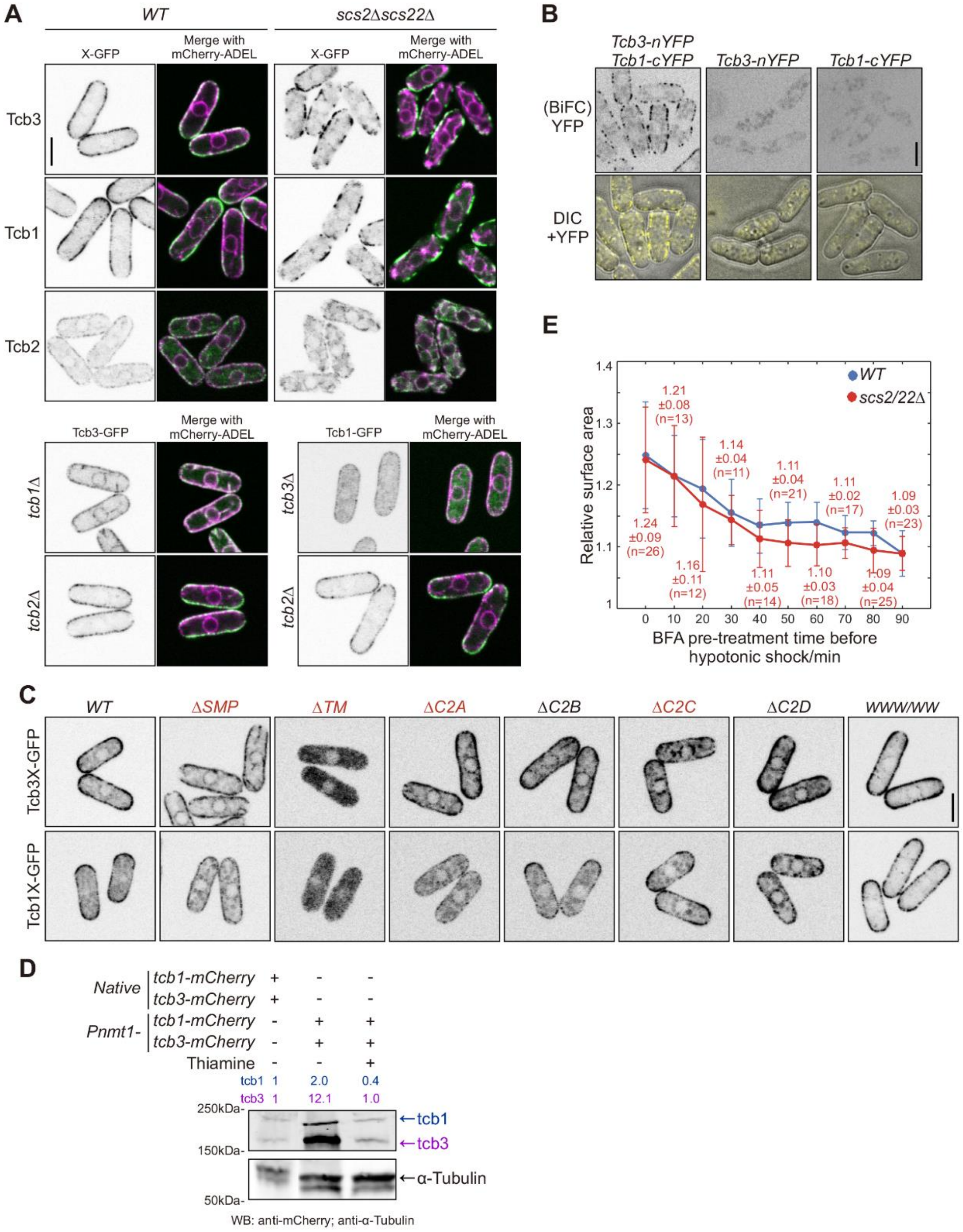
A
WT
scs2Δscs22Δ
X-GFP
Merge with mCherry-ADEL
Tcb3
Tcb1
Tcb2
Tcb3-GFP
Tcb1-GFP
tcb1Δ
tcb2Δ
tcb3Δ
tcb2Δ
B
Tcb3-nYFP Tcb1-cYFP
Tcb3-nYFP
Tcb1-cYFP
(BiFC) YFP
DIC +YFP
E
WT
scs2/22Δ
Relative surface area
BFA pre-treatment time before hypotonic shock/min
1.24 ±0.09 (n=26)
1.21 ±0.08 (n=13)
1.16 ±0.11 (n=12)
1.14 ±0.04 (n=11)
1.11 ±0.05 (n=14)
1.11 ±0.04 (n=21)
1.10 ±0.03 (n=18)
1.11 ±0.02 (n=17)
1.09 ±0.04 (n=25)
1.09 ±0.03 (n=23)
C
WT
ΔSMP
ΔTM
ΔC2A
ΔC2B
ΔC2C
ΔC2D
WWW/WW
Tcb3X-GFP
Tcb1X-GFP
D
Native
tcb1-mCherry
tcb3-mCherry
Pnmt1-
Thiamine
tcb1 1 2.0 0.4
tcb3 1 12.1 1.0
250kDa-
150kDa-
50kDa-
←tcb1
←tcb3
←α-Tubulin
WB: anti-mCherry; anti-α-Tubulin

**Figure S3 Tcb1-Tcb3 complex mediates lipid transfer at ER-PM contacts.**

(A and C) Central focal plane scanning confocal images of indicated cells. (B) Central focal plane spinning disk confocal images of indicated cells. Scale bar, 5 µm. (D) Western blot of indicated proteins expressed from native loci or under *nmt1* promoter. Samples were probed with indicated antibodies. Thiamine, suppressing expression of nmt1 promoter. Quantification, relative protein expression level. (E) Quantifications of maximum relative surface area (mean ± SD shown for *scs2Δscs22Δ*) of BFA-pretreated *WT* and *scs2Δscs22Δ* protoplasts after 10-min acute hypotonic shock. The relative surface area of *WT* at each time points are 1.25±0.8 [n=26], 1.23±0.7 [n=13], 1.17±0.8 [n=12], 1.16±0.5 [n=11], 1.15±0.5 [n=14], 1.15±0.6 [n=21], 1.14±0.4 [n=18], 1.12±0.4 [n=17], 1.12±0.4 [n=25], and 1.09±0.4 [n=23] (mean ± SD). n, cell number; Error bars, 2×SD.

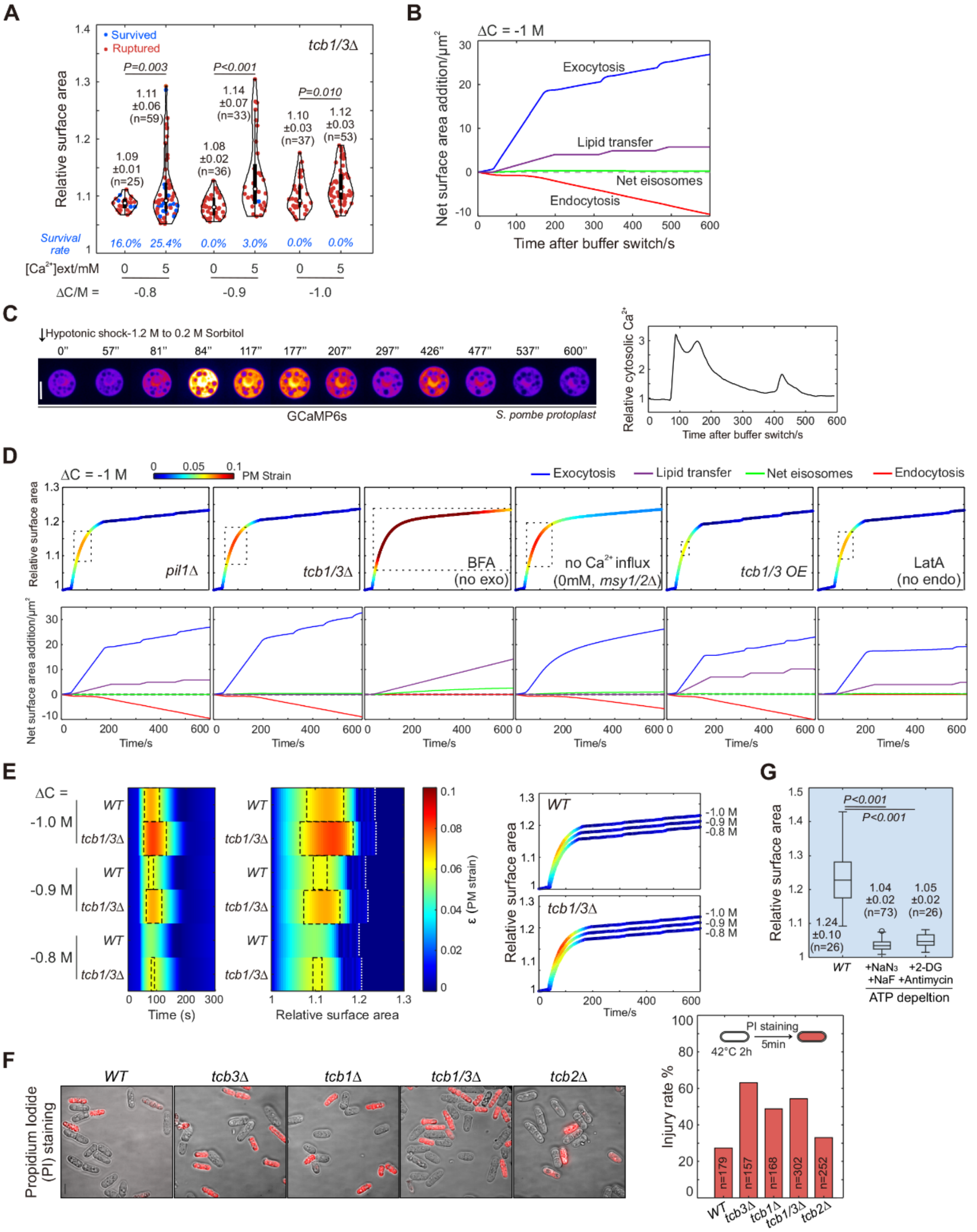
A
Survived
Ruptured
tcb1/3Δ
Relative surface area
P=0.003
P<0.001
P=0.010
1.09 ±0.01 (n=25)
1.11 ±0.06 (n=59)
1.08 ±0.02 (n=36)
1.14 ±0.07 (n=33)
1.10 ±0.03 (n=37)
1.12 ±0.03 (n=53)
Survival rate
16.0%
25.4%
0.0%
3.0%
0.0%
0.0%
[Ca2+]ext/mM
ΔC/M =
-0.8
-0.9
-1.0
B
ΔC = -1 M
Net surface area addition/μm2
Exocytosis
Lipid transfer
Net eisosomes
Endocytosis
Time after buffer switch/s
C
Hypotonic shock-1.2 M to 0.2 M Sorbitol
0'' 57'' 81'' 84'' 117'' 177'' 207'' 297'' 426'' 477'' 537'' 600''
GCaMP6s
S. pombe protoplast
Relative cytosolic Ca2+
Time after buffer switch/s
D
ΔC = -1 M
PM Strain
Exocytosis
Lipid transfer
Net eisosomes
Endocytosis
Relative surface area
pil1Δ
tcb1/3Δ
BFA (no exo)
no Ca2+ influx (0mM, msy1/2Δ)
tcb1/3 OE
LatA (no endo)
Net surface area addition/μm2
Time/s
E
ΔC =
-1.0 M
-0.9 M
-0.8 M
WT
tcb1/3Δ
Time (s)
Relative surface area
ε (PM strain)
Time/s
G
P<0.001
P<0.001
Relative surface area
1.24 ±0.10 (n=26)
1.04 ±0.02 (n=73)
1.05 ±0.02 (n=26)
WT
+NaN3 +NaF
+2-DG +Antimycin
ATP depeltion
F
WT
tcb3Δ
tcb1Δ
tcb1/3Δ
tcb2Δ
Propidium Iodide (PI) staining
PI staining
42°C 2h
5min
Injury rate %
n=179
n=157
n=168
n=302
n=252

**Figure S4 A numerical model describes the hypotonic PM expansion under different hypoosmotic shocks.**

(A) Quantifications of maximum relative surface area (mean ± SD) of indicated protoplasts after 10-min acute hypotonic shock of indicated conditions. (B) Net surface area added by each source during hypoosmotic expansion within 10 minutes, for same model parameters as Figure 4B. (C) Time-lapse spinning disk confocal images of representative protoplasts expressing GCaMP6s with indicated hypotonic shocks. Shown are pseudo-colored images at central focal planes. Times, relative to buffer switch. Graph shows r$Ca^{2+}$Cyto curves of the representative cell. (D) Simulation of relative PM surface area expansion in indicated scenarios with colors indicating PM strains (ε) and net surface area added by each source during 10-min hypotonic expansion. The framed vulnerable period is defined as ε above 0.06. (E) Plots of simulated ε variation in indicated model conditions along hypoosmotic expansion time (left) and relative surface area (middle). Vulnerable periods are framed. Dashed lines, the maximum relative surface area after reaching homeostasis. Simulation of relative PM surface area expansion (right) in indicated scenarios. ε is indicated by colors. (F) Spinning disk confocal image of PI-stained walled cells. Injury rate is defined by the percentage of cells stained by PI. Scale bar, 5 µm. (G) Box-and-whisker plot of maximum relative surface area (mean ± SD) of indicated protoplasts after 10-min acute hypotonic shock of indicated conditions. n, cell number; *P*-values, two-tailed t-test.
